# Spotlight on 10x Visium: a multi-sample protocol comparison of spatial technologies

**DOI:** 10.1101/2024.03.13.584910

**Authors:** Mei R. M. Du, Changqing Wang, Charity W. Law, Daniela Amann-Zalcenstein, Casey J. A. Anttila, Ling Ling, Peter F. Hickey, Callum J. Sargeant, Yunshun Chen, Lisa J. Ioannidis, Pradeep Rajasekhar, Raymond K. H. Yip, Kelly L. Rogers, Diana S. Hansen, Rory Bowden, Matthew E. Ritchie

**Author notes:** These authors contributed equally as co-first (Mei R. M. Du, Changqing Wang, Charity W. Law, Daniela Amann-Zalcenstein). These authors contributed equally as co-last (Diana S. Hansen, Rory Bowden, Matthew E. Ritchie).

## Abstract

**Background:** Spatial transcriptomics allows gene expression to be measured within complex tissue contexts. Among the array of spatial capture technologies available is 10x Genomics’ Visium platform, a popular method which enables transcriptome-wide profiling of tissue sections. Visium offers a range of sample handling and library construction methods which introduces a need for benchmarking to compare data quality and assess how well the technology can recover expected tissue features and biological signatures.

**Results:** Here we present *SpatialBench*, a unique reference dataset generated from spleen tissue of mice responding to malaria infection spanning several tissue preparation protocols (both fresh frozen and FFPE samples, with and without CytAssist tissue placement). We noted better quality control metrics in reference samples prepared using probe-based capture methods, particularly those processed with CytAssist, validating the improvement in data quality produced with the platform. Our analysis of replicate samples extends to explore spatially variable gene detection, the outcomes of clustering and cell deconvolution using matched single-cell RNA-sequencing data and publicly available reference data to identify cell types and tissue regions expected in the spleen. Multi-sample differential expression analysis recovered known gene signatures related to biological sex or gene knockout.

**Conclusions:** We framed a comprehensive multi-sample analysis workflow that allowed us to generate consistent results both within and between different subsets of replicate samples, enabling broader comparisons and interpretations to be made at the group-level. Our *SpatialBench* dataset, analysis, and workflow can serve as a practical guide for Visium users and may prove valuable in other benchmarking studies.

## Background

Spatial transcriptomic technologies allow gene expression to be measured in complex tissue samples in an x-y context (1, 2). The main approaches for spatially resolving transcript expression rely on either imaging based in-situ hybridization-based methods (e.g. MERFISH (3), seqFISH (4) and CosMx SMI (5)), in-situ sequencing-based methods (e.g. STARmap (6) and HybISS (7)) or array-based sequencing protocols (e.g. Visium (8), Slide-Seq1&2 (9, 10) and Stereo-Seq (11)).

Methods vary considerably in terms of the spatial resolution they allow (from sub-cellular to multi-cell) and number of features that can be measured (from small focused gene panels to genome-wide expression). The rapid expansion in both the spatial transcriptomic protocols available to researchers and subsequent analysis methods (12) introduces a need for benchmarking to compare the performance of different combinations of platforms and analysis approaches (13). Recent cross-platform benchmarking efforts include the *cadasSTre* project (14) which focuses on sequencing-based methods across a range of mouse tissues, comparison analyses of imaging-based methods on human cancer and mouse brain tissue (15, 16), and *SpaceTx* which includes both imaging and sequencing-based technologies using human brain and mouse primary visual cortex tissue (17). Other benchmarking studies aim to evaluate the performance of analysis methods developed for different pre-processing and downstream procedures (18–21).

By far the most popular sequencing-based method at present is the commercially available Visium method from 10x Genomics. Visium’s experimental process involves the capture of spatially barcoded mRNA transcripts on a slide, followed by reverse transcription, library preparation and sequencing. The resulting data integrates gene expression profiles with spatial coordinates. A notable feature is Visium’s versatility in terms of sample compatibility, being able to accommodate both fresh frozen (OCT) and formalin-fixed, paraffin-embedded (FFPE) samples, expanding its applicability to a broad range of tissue types and experimental conditions. This flexibility allows researchers to leverage existing FFPE archives, overcoming the limitation of previous single-cell and spatial technologies that are restricted to OCT preserved tissue (22).

A question that requires exploration for Visium technology was how different sample handling methods affect data quality, spatially variable gene detection and downstream analysis results. To address this, we generated the *SpatialBench* reference dataset, which includes replicate tissue sections that span different sample handling methods, including fresh frozen with manual tissue sectioning and polyA library preparation and the CytAssist (CA) automated method that uses a probe-based protocol, as well as FFPE with manual sectioning or CytAssist (which both use probe-based protocols).

Mouse spleens responding to malaria infection were selected as a reference tissue in our study. As *Plasmodium spp*. are blood-borne parasites, the spleen constitutes a key site in the immune response to infection, with antibody responses playing an important role in protection (23–27). The development of this antibody-mediated immunity requires the establishment of germinal centre (GC) structures in lymphoid organs, where activated B cells undergo antibody affinity maturation. GC responses to malaria have been found to be regulated by transcription factors, such as *T-bet*, which are preferentially activated in response to the highly inflammatory milieu elicited during acute infection (28, 29). This infection thus provides an excellent experimental system to not only investigate functional organ architecture but also analyse specific structures within the spleen only visibly upregulated in response to an active infection. Samples from both male and female mice along with *Tbx21*^*fl/fl*^*Cd23*^*Cre*^ (conditional knockout of T-bet in mature follicular B cells) samples were included in our study, allowing us to explore our ability to recover sex-specific gene signatures as well as examine spatial influence of *T-bet* on GC response.

We use these data to compare sample handling and analysis methods, develop a workflow for multi-sample analysis that is able to recover expected ground truth in terms of both tissue architecture and differential expression of known gene signatures.

## Results

### *SpatialBench* reference samples profiled using 10x Visium and scRNA-seq technology across different sample handling *×* protocol combinations

Our study profiled mouse spleen responding to malaria infection (Figure 1a) in a total of 13 samples (Figure 1b) sequenced across 4 experiments on an Illumina NextSeq 2000 instrument (Supplementary Table S1). The samples were processed in 4 different ways (or “Sample Types”): OCT, OCT CA, FFPE, and FFPE CA (Figure 1b). These refer to tissue preservation, either as fresh frozen at optimal cutting temperature (OCT) or formalin-fixed paraffin-embedded (FFPE), and tissue placement, either directly placed (Manual) or using CytAssist (CA). Samples were of different genotypes and sexes (Wild Type (WT) females, T-bet knockout (KO male) or control (CTL male), where the KO and CTL samples were available as OCT preserved only. We grouped these samples by sample type and will refer to them broadly as: FFPE Experiment 1, OCT Experiment 2, FFPE Experiment 3, and CA Experiment 4 (Supplementary Table S1). Samples processed by FFPE and also those prepared with CytAssist use probe-based ligation protocols for library construction, whereas OCT with manual tissue placement uses a poly-Abased capture method. A separate matching single-cell RNAseq dataset was also generated from three FFPE samples.

**Fig. 1.**
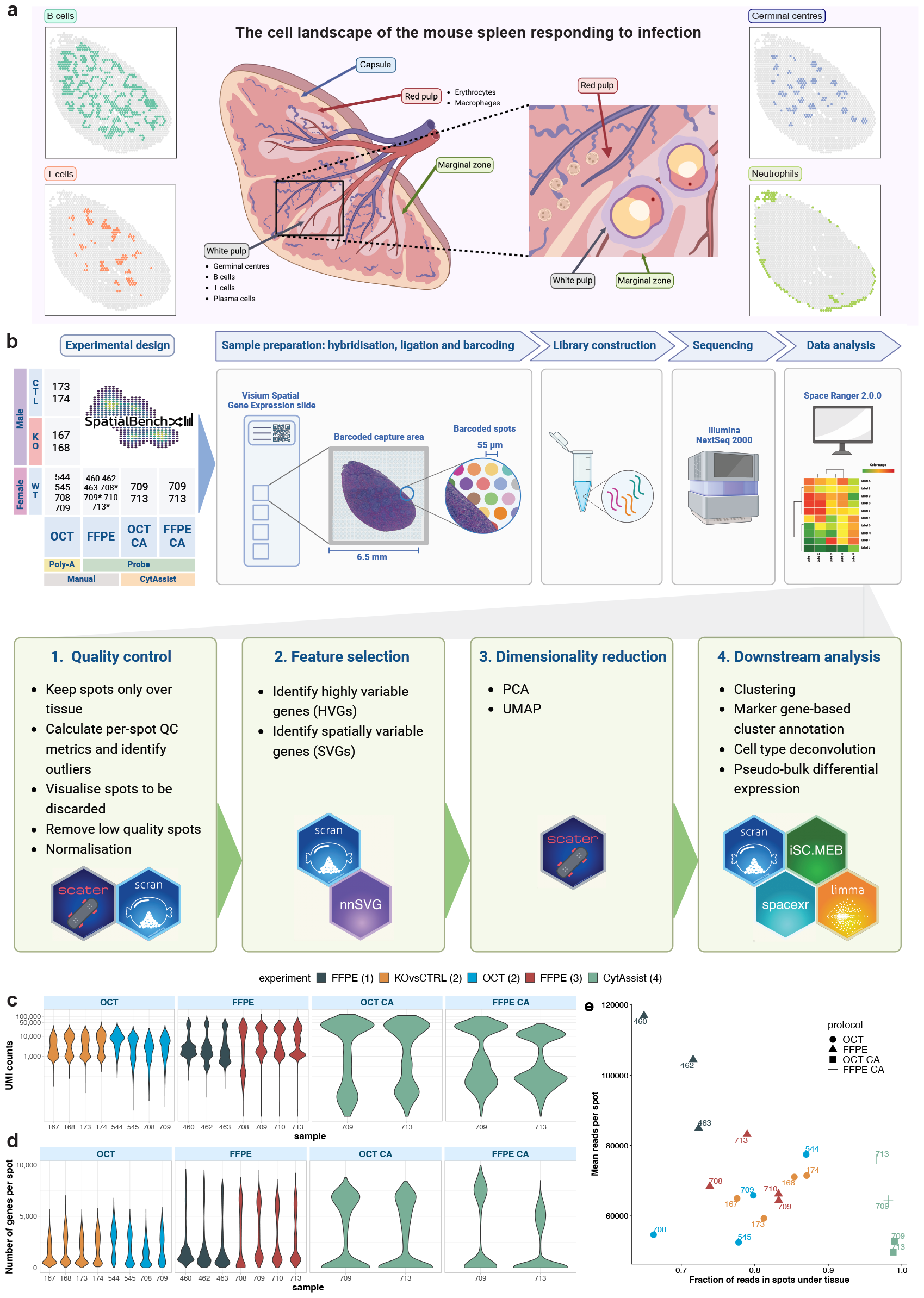
Overview of the experimental workflow, data generated and its analysis. (**a**) The mouse spleen and major cell types (B, T and Plasma cells, Erythrocytes, Neutrophils and Macrophages) and structures (Germinal centres which are predominantly made up of B cells) expected following infection, which are organised into broader tissue regions (Red and White pulp). Figure created with BioRender.com. (**b**) 13 samples were captured over 4 10x Genomics Visium OCT slides and 3 FFPE slides, and sequenced over 5 runs on an Illumina NextSeq 2000. Samples are categorised by sex, genotype, tissue preparation protocol, library construction protocol, and tissue placement. A matching scRNA-seq sample of 3 mouse spleens (samples denoted with an asterisk) was captured over 1 gel bead-in emulsion (GEM) well, and gene expression and hashtag oligos (HTOs) were sequenced over 1 Illumina run. Subsequent data analysis involved processing with 10x Genomics Space Ranger 2.0.0, and quality control, feature selection, dimensionality reduction and downstream analysis using various R-based software packages. Figure created with BioRender.com. (**c**) Violin plots of UMI counts per spot for all samples, grouped by tissue preparation protocol. The y-axis is on a log_10_ scale for clarity. (**d**) Violin plots of number of genes detected per spot for all samples, grouped by tissue preparation protocol. (**e**) A scatterplot showing the fraction of reads captured by spots under tissue against the mean number of reads per spot. The order of experiments is reflected in the shared legend.

The data processing workflow builds upon scRNA-seq analysis, with the additional use of spatial coordinate information in steps such as feature selection and clustering (Figure 1b). An overview of our dataset shows that across all experiments, probe-based samples had higher UMI counts and numbers of detected genes, particularly the CytAssist samples (Figure 1c, d). Spots located beyond tissue boundaries, and many spots with low UMI counts and few detected genes were filtered out during quality control (Figure 1b).

### Probe-based samples have higher UMI counts

Tissue sections were placed on a Visium slide containing 4,992 spots that were used to measure gene expression. The number of spots that were covered by tissue depends on both its size and shape. Across our experiments using mouse spleen, an average of 1,957 spots (39%) were covered by tissue (range: 592-3,224, see Supplementary Table S2). The amount of sequencing carried out for each sample was adjusted such that larger tissue sections were sequenced more deeply, to have on average the same number of reads per spot than smaller tissue samples.

Samples had a mean of 39,616,270 valid UMI counts, that is, UMIs covered by tissue. Across each experiment, polyA-based OCT Experiment 2 had a mean of 23,642,694 valid UMI counts, while for probe-based experiments (FFPE Experiment 1 and 3, CA Experiment 4) this was higher at 41,630,649. Separately, CA Experiment 4 had the highest mean valid UMI count of 70,815,948, and FFPE Experiment 3’s mean (50,309,451) was almost double that of FFPE Experiment 1 (26,355,327).

Poly-A-based OCT samples had a mean of median UMI counts per tissue-covered spot of 8,360. This is lower than our probe-based experiments which had higher sensitivity; with FFPE Experiment 1 having a median of 33,390 UMI counts per spot, FFPE Experiment 3 at 21,730, and CA Experiment 4 at 24,804. Within each experiment, UMI counts were generally consistent across samples. FFPE CA sample 713, however, had a notably lower UMI count and a median UMI count of less than half compared to the other CA samples (Figure 2a). Differences between poly-A and probe-based experiments are shown for sample 709 as an example in Figure 2b and c. They can also be observed in OCT samples (KO vs CTL and OCT) having lower counts in Supplementary Figure S1. An edge bias was also apparent, characterised by higher UMI counts along the edges of the OCT samples. This raises the possibility of under-permeabilisation following tissue optimisation, before library preparation and sequencing.

**Fig. 2.**
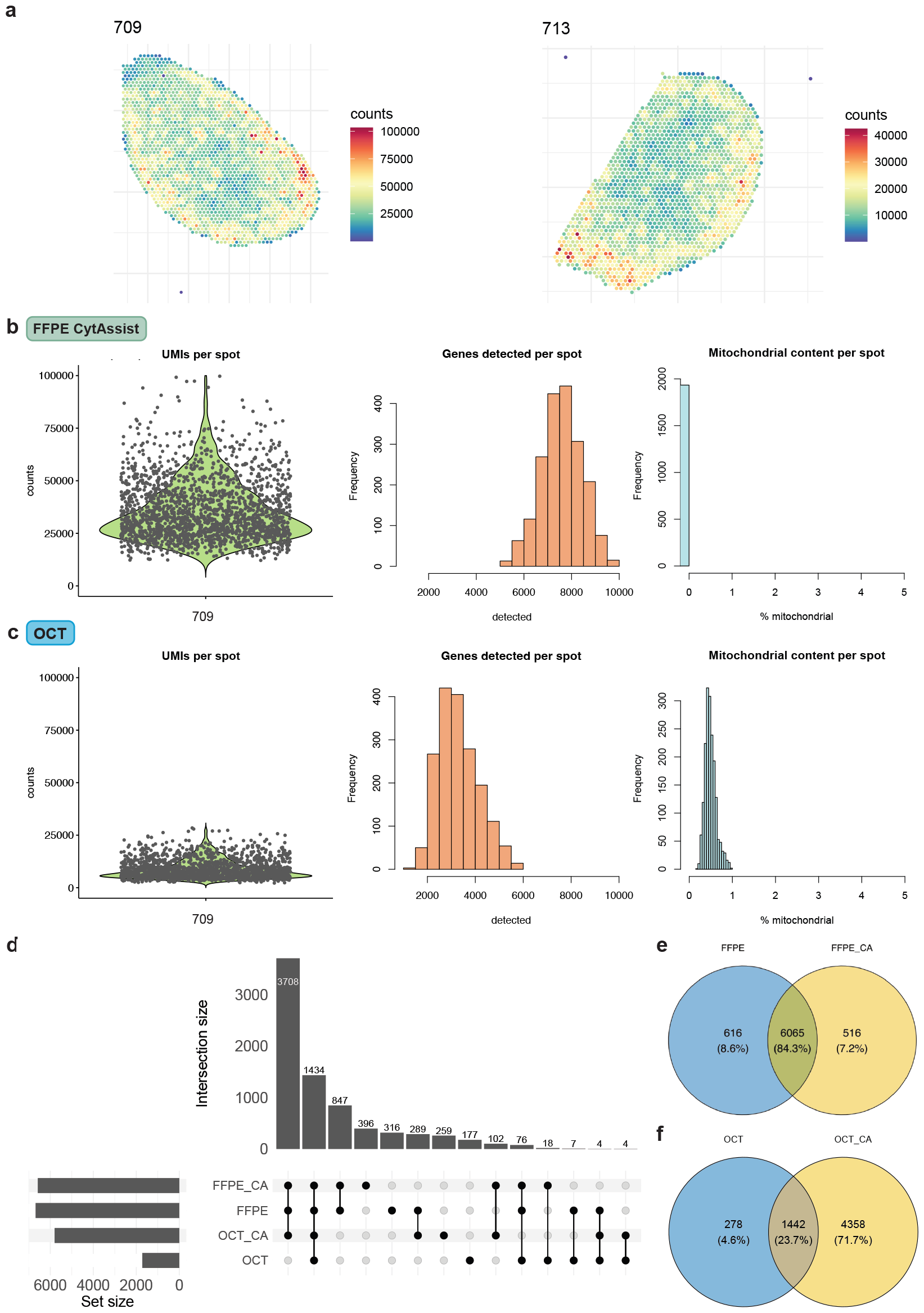
Quality control procedures. (**a**) The spatial distribution of UMI counts per spot in FFPE CytAssist (CA) samples 709 and 713. (**b**) Quality control metrics for FFPE CA sample 709 following filtering with *scater*. (**c**) Quality control metrics for OCT sample 709 following filtering with *scater*. (**d**) UpSet plot showing the overlap of detected genes in all wild type (WT) samples, categorised by sample type. Detected genes are defined as genes with a count of 3 or more in at least 10% of spots. (**e**) Venn diagram of genes in FFPE samples, with and without CA. (**f**) Venn diagram of genes in OCT samples, with and without CA.

### Probe-based samples have higher mapping confidence

Reads mapped with more than 85% confidence to the probe set in our probe-based experiments. All samples in FFPE Experiment 1 and CA Experiment 4 had reads mapped with more than 97% confidence, whilst FFPE Experiment 3 ranged between 85-98%. This was expected as probe sets are highly specific to known sequences in the reference transcriptome. We also observed that poly-A-based experiments (Experiment 2 plus two related OCT experiments not included in this dataset) had lower values of reads being mapped confidently to the transcriptome (66-79%).

### CytAssist facilitates the capture of more reads under tissue

Importantly, although some samples from earlier experiments had higher sequencing read numbers, the fraction of reads in spots under tissue was close to 100% for samples placed with CytAssist (Figure 1e). This is another indication of higher quality overall, as a lower proportion of reads would have been filtered out from these samples during quality control. In contrast, approximately 65-87% of reads fell within tissue boundaries for OCT and FFPE samples. For some samples like OCT 460 and FFPE 708, a notable amount of sequencing reads were assigned to spots outside the boundary of the tissue sections. Tissue boundaries are annotated by imaging processing software, but some images or areas of the tissue can be blurry and boundary definitions can be inaccurate, making it difficult to decide whether a spot falls within or outside of tissue boundaries. The improvement in reads captured within tissue boundaries for CytAssist experiments may be attributed to enhancements implemented in the CytAssist platform.

### Removing low quality spots

The initial phase of all analysis workflows involves evaluating data quality. This includes the detection of outlier spots, removal of low-quality spots, and normalisation (Figure 1b). Only spots covered by tissue were retained. They were then further filtered out from downstream analyses if specific criteria for library size, detected features and mitochondrial content were not met, indicative of low quality. Per-spot metrics were computed, and outliers were identified using the *scater* (30) package. Following spot filtering, normalisation using scRNA-seq methods from *scater* and *scran* (31) were applied.

Following quality control, the proportion of spots covered by tissue remaining for further analysis was greater than 0.90 for all probe-based samples, except FFPE sample 708 (0.87) (Supplementary Table S2). On average, this was 0.95 for probe-based samples, compared to 0.83 for poly-A-based samples, as lower proportions of UMIs and spots were removed, especially in CA samples. This led to higher proportions of UMIs per spot and spots under tissue passing quality control and being retained for downstream analysis. FFPE Experiment 1 samples retained an even higher average proportion of spots (0.96) following filtering. However, these samples initially started with low numbers (*<*1,000) and low proportions (on average, 0.14) of spots under tissue. Additionally, they displayed the highest proportions of UMIs removed among probe-based samples, further restricting the number of spots available for downstream analysis.

### Different genes are detected for poly-A-based OCT samples than other sample types

The poly-A-based capture method, used for OCT samples, selects genes or transcripts by their poly-A tail, in theory allowing any expressed gene to be detected given sufficient sequencing. This is in contrast with the other sample types (OCT CA, FFPE and FFPE CA), where genes are selected using a uniform set of probes. For these probe-based samples, only genes that are both expressed and within the probe set can be detected. However, as the number of features is smaller in the probe set, more genes can be detected at a given sequencing level. Across our samples, we identified 1,636 genes detected in OCT samples that used poly-A-based gene selection (Figure 2d). Here, we define detected genes as genes with a count of 3 or more in at least 10% of spots under tissue remaining after quality control. For probe-based gene selection, a much higher number of genes were detected; 5,800 genes for OCT CA samples and 6,681 for FFPE and 6,581 for FFPE CA samples. Sample 709 is used to show these differences in Figure 2b and c. In total, 38% of 19,465 genes targeted in Visium Mouse Transcriptome Probe Set v1.0 were detected in the probe-based samples. The largest overlap in detected genes is between the 3 probe-based sample types. The effect of probe-based gene selection on OCT tissue for detected genes is apparent in Figure 2f, as there is a much smaller overlap (22.6%) between OCT and OCT CA samples, compared to the high overlap (84.3%) between FFPE and FFPE CA samples.

At a more lenient threshold of detected genes being defined as genes with a count of 3 or more in at least 1% of spots (Supplementary Figure S2), the largest overlap in detected genes instead includes all 4 sample types. This implies that most of the original overlapping genes between poly-A-based and probe-based samples were very lowly expressed under the poly-A protocol. Similar numbers in the overlap of probe-based samples were observed for both thresholds, 4,000 genes.

It is worth noting that specific sets of genes, such as mitochondrial genes, were missing from probe-based samples (Figure 2b), making detected genes inconsistent between poly-A-based and probe-based samples. Additionally, there were 177 genes detected only in OCT samples (Figure 2d), of which there were no mitochondrial genes, but 57% were either ribosomal or mitochondrial ribosomal protein genes. These were also not detected in the probe set as expected. However, of the remaining 43%, more than half were included in the probe set, although not detected in the probe-based samples according to the detection threshold used. These consisted mainly of mitochondria-related genes, though a few immune-related genes were detected such as *Csf1*, characteristic of red pulp macrophages (32), and *Cd7* expressed in T cells (33). OCT-only genes that were not found in the probe set include *Hba-a1* expressed in erythrocytes (34), and *Ccl19* and *Ccl21a* involved in T cell immune responses (35).

### Downstream analysis by sample type

Following preprocessing of individual samples, we applied a standard Bioconductor workflow (36) to explore feature selection and clustering. Highly variable genes (HVGs) could be identified using established scRNA-seq methods from the *scran* package (31) (Figure 3a, top row), though HVGs are notably defined based on expression data along and do not consider spatial information. Clustering was also possible following scRNA-seq methods with default parameters, deriving clusters in each sample without using spatial coordinates. However, different numbers of clusters were obtained for each sample (Figure 3b).

**Fig. 3.**
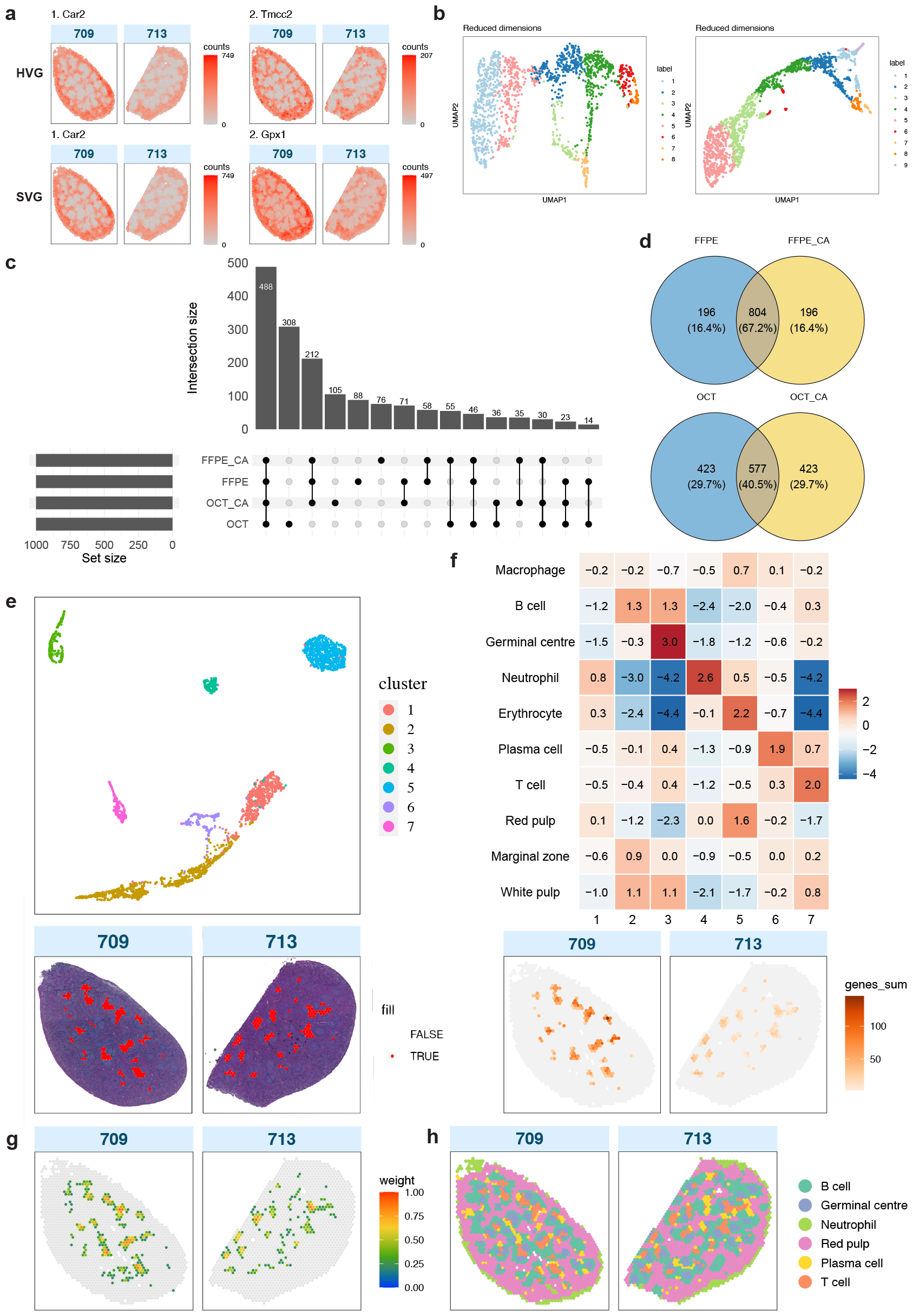
Summary of downstream analyses. (**a**) Top: Spatial expression of the top 2 HVGs for FFPE CA samples. HVGs were identified for each sample, but are shown together here, as the top 2 HVGs were the same in both samples. Bottom: Spatial expression of the top 2 SVGs for FFPE CA samples. These were identified in a single gene list generated through a multi-sample approach. (**b**) Clusters identified in each FFPE CA sample following a standard Bioconductor workflow. (**c**) UpSet plot showing the overlap of the top 1,000 SVGs in each sample type for all wild type (WT) samples. (**d**) Venn diagrams showing unique and overlapping SVGs between FFPE samples and between OCT samples, with and without CytAssist, among the top 1,000 SVGs. (**e**) Top: A UMAP plot showing spatial clusters across both FFPE CA samples, identified by *iSC*.*MEB*. Bottom: Spatial cluster 7 projected onto tissue images. (**f**) Top: A heatmap showing scores for expression of cell type marker gene groups in each cluster compared to all other clusters in FFPE CA samples. Bottom: The aggregate gene expression of the T cell marker gene group in cluster 7. (**g**) A deconvolution plot of confident weights for T cells generated by *spacexr* for FFPE CA samples. (**h**) FFPE CA samples 709 and 713 with each spot annotated following deconvolution and marker gene expression analysis, using spatial clusters.

In addition to not using spatial information, a limitation of using common feature selection and clustering methods was the necessity to process samples individually, making it difficult to derive broader insights between both biological and technical replicate samples. This challenge is further complicated when comparing samples from different conditions, such as KO and CTL genotypes. For example, Cluster 1 in one sample may not correspond to Cluster 1 in another sample, necessitating the need for labels to be resolved through cell type deconvolution. For a more integrative analysis approach, we processed multiple samples simultaneously by sample type, allowing for consistent cluster labels within each batch.

### Feature selection

We identified spatially variable genes (SVGs) using *nnSVG* (37), making use of the spatial information in our data (Figure 3a, bottom row). *nnSVG* was run in multi-sample mode, which firstly finds SVGs in individual samples and ranks them. Ranks of genes that reached statistical significance with an adjusted *p*-value below 0.05 are averaged across all the samples in a batch to produce a cross-sample rank. Additional gene filtering was performed individually per sample and log *−* counts were then re-calculated. The output is a single list of SVGs and their overall ranks across all replicates of the same sample type. When comparing the top-ranked (1,000) SVGs between different sample types, we observe that, reassuringly, the largest overlap are genes shared in common between all 4 sample handling methods (Figure 3c). Within the FFPE and OCT groups, there is also relatively high agreement between FFPE and FFPE CA samples, and OCT and OCT CA samples have equal proportions of unique SVGs (Figure 3d). The second largest category corresponds to OCT-specific SVGs, drawing attention to the systematic differences between poly-A and probe-based results highlighted previously. However, upon considering all significant SVGs, we find more genes are identified as spatially variable in probe-based platforms, with 3,380 SVGs from the OCT samples versus a mean of 8,555 genes for probe-based sample types (OCT CA, FFPE and FFPE CA) (Supplementary Figure S3a). This observation is also supported by the higher intersection of FFPE and FFPE CA SVGs, and OCT CA only genes occupying a greater percentage of all OCT SVGs (Supplementary Figure S3b). These findings reinforce the need to analyse OCT samples that used poly-A-based gene selection separately from samples that use the probe-based version of the Visium technology.

There is a high degree of overlap between the top SVGs and top HVGs, indicating that a significant amount of biological signal in our dataset is captured by the spatial distribution of cells, particularly in red pulp and white pulp. For FFPE CA samples, the highest overlap between all gene lists HVGs from sample 709, HVGs from sample 713 and multi-sample FFPE CA SVGs was *∼* 0.76, at the intersection of 537 genes (Supplementary Figure S3c). These lists also share the same top gene, *Car2* (Figure 3a), following removal of highly abundant haemoglobin and immunoglobulin genes (38).

Gene Ontology (GO) terms enriched amongst the top 1,000 SVGs in FFPE CA samples relate to immune activation and regulation, with focus on leukocytes and lymphocytes like B cells, which are active during an immune response (39, 40) (Supplementary Figure S3d). Inspecting the spatial distribution of additional highly ranked SVGs reveals two distinct cell clusters and gene expression profiles (Supplementary Figure S4). It is clear from pathology image analysis using a trained classifier (Supplementary Figure S5) and further downstream analysis that these reflect the distribution of red pulp and white pulp, the two major regions of the spleen (41).

### Clustering

Methods adapted from single-cell analysis workflows were previously demonstrated on individual samples. In our multi-sample approach with spatially aware clustering, we could derive clusters that were concordant across all samples types using the *iSC*.*MEB* (42) package. Normalised counts of each sample in a sample type group were combined into a single matrix for *iSC*.*MEB* to perform principal component analysis (PCA) to obtain principal components (PCs). The top 10 PCs were then used for spatial clustering to obtain cluster labels. *iSC*.*MEB* also offers differential expression analysis, which was used in guiding cluster refinement to avoid over-clustering of spots. As a result, we identified 7 spatial clusters in FFPE CA samples (Figure 3e), and both samples appear to have been integrated evenly following batch correction (Supplementary Figure S6).

### Cell type deconvolution

The high-performing *spacexr* (21, 43) (formerly Robust Cell Type Decomposition (RCTD)), was chosen to perform cell type deconvolution. *spacexr* uses annotated scRNA-seq data to generate gene expression profiles for each cell type in the reference. It then fits a probabilistic model to estimate cell type proportions for each spot and creates spatial maps of cell types, fitting each spot as a linear combination of individual cell types. There are different modes to process data with one, two or an unknown number of cells in each spot. We used the latter multi-mode recommended for Visium, which can accommodate for more than two cell types per spot and considers all such cell types when estimating proportions. We also used our matching scRNA-seq reference dataset. Estimates can be classified as confident, and the distribution of confident weights inferring cell type proportion can be visualised in a spatial map as shown in Figure 3g. From this analysis, we were able to consolidate the classification of major cell types B cells, T cells, erythrocytes, and neutrophils.

To evaluate the performance of our matching scRNA-seq reference, we also performed deconvolution with a public mouse spleen dataset (44). We compared the proportion of spots with confident assignments as a metric, defining confidence as a confident weight calculated by *spacexr* of greater than 0.5. The public reference was more finely annotated initially and required additional grouping of cell types into broader categories, after which it produced similar results to those generated with our matching reference.

With the public reference, proportions ranged from 0.73-0.86 for FFPE samples, 0.60-0.90 for OCT samples, 0.85-0.89 for FFPE CA samples, and 0.83-0.93 for OCT CA samples. With our matching reference, proportions ranged from 0.65-0.85 for FFPE samples, 0.49-0.91 for OCT samples, 0.83-0.84 for FFPE CA samples, and 0.88-0.94 for OCT CA samples. Samples in FFPE Experiment 1 (460, 463, and 463) were also observed to have lower proportions (0.65-0.79) than those in FFPE Experiment 3 (0.80-0.85) when using the *SpatialBench* reference. Notably, all CA samples had high proportions above 0.8, though overall, these results are not necessarily indicative of data quality. For example, FFPE sample 709 has more mean reads per spot and a higher fraction of reads in spots under tissue than OCT sample 708 (Figure 1e), but its proportion metric is lower at 0.81, compared to OCT sample 708’s 0.91.

We also examined the expression of marker genes selected from previous studies and literature search (see Methods) that are particular to the cell types (B, T and plasma cells, macrophages, neutrophils and erythrocytes) expected in a spleen responding to infection. We developed a cluster score approach using groups of marker genes that could be used to identify a cell type or tissue region (Supplementary Table S3), defining the score as the log-fold change between one cluster and all other clusters (see Methods). This is demonstrated in Figure 3f for FFPE CA samples, which shows distinct scores reflecting prominent cell types, for clusters 3-7 in particular. The score for (red pulp) macrophages was not as high as for red pulp and erythocytes, which may be due to the exclusion of some marker genes from the probe set. Cluster scores for the major regions present in this spleen model (germinal centre, red pulp, marginal zone and white pulp) were also included to investigate their cell type compositions. Their scores signify that mostly erythrocytes constitute the red pulp, B cells and some T cells can be seen in the marginal zone, and germinal centres, B cells and T cells are located in white pulp, aligning with their expected distributions (45). The strong signal for germinal centres also reflects the presence and robust activity of these structures during an immune response.

### Spatial cluster annotation

We combined our results from all the above steps in the multi-sample downstream analysis to assign cell type labels to each spatial cluster. It is important to consider that Visium does not provide single-cell resolution, and therefore a singular label is not entirely reflective of the cell type proportions at each spot. Nonetheless, we assigned labels for the most abundant cell type in each cluster and showcase an effective visual representation of the mouse spleen during infection (Figure 3h).

While the classification of most cell types was relatively straightforward, varied results were presented during the assignment of red pulp clusters. This issue was observed across all sample type groups. The top HVG and SVG in FFPE CA samples, *Car2*, is a marker gene of erythrocytes (46) and can be seen expressed in a distinctive spatial pattern, with a wide outer edge around the spleen and in a series of circular structures in the middle of the tissue (Figure 3c). The distribution of confident weights for erythrocytes exhibited a similar pattern following cell type deconvolution (Supplementary Figure S7). Our image segmentation classifier (see Methods) also mirrored this pattern, showing a clear distinction between the red pulp and white pulp regions (Supplementary Figure S5). However, the clusters in question, 1 and 5, appeared to be distinct in the cluster UMAP (Figure 3e) and cluster scores (Figure 3f), separating into inner and outer regions of spots. Yet, the highest score for cluster 1, albeit low, pointed to erythrocytes. Ultimately, these clusters were both labelled as red pulp.

Overall, our results align with the the structures and lack of architecture consistent with malaria infection (45) (Figure 1a). There is less organisation and definition observed in the structures containing B cells and T cells, B cell follicles and T cell zones respectively, and transient loss of marginal zones (Figure 3h). A key detail is the striking presence of germinal centres, which are formed in response to infection.

### Comparing gene expression between replicate male and female spleens

We also used sex-specific differences between samples as ground truth to evaluate differential expression between CTL (male) and WT (female) OCT samples from Experiment 2. We clustered spots jointly for all CTL and WT OCT samples (Figure 4a and Supplementary Figure S8), identifying red pulp (clusters 1, 4 and 5), B cell (cluster 2), neutrophil (cluster 3), germinal centre (cluster 6), plasma cell (cluster 7) and T cell (cluster 8) enriched spots using the marker genes described above (Figure 4b and c). For each cluster, we then aggregated spot counts at the sample-level, filtered genes with low counts using *edgeR* (47, 48) and performed differential expression (DE) analysis with the *limmavoom* (49) pipeline comparing male versus female samples. Figure 4d shows an MA plot (log-fold-change versus average expression) for the male versus female DE comparison within the B cell cluster, with plots for the other clusters included in Supplementary Figure S9. Chromosome X genes that were previously identified as X inactivation escape genes in mouse spleens (50) (17 genes were in the original list, one of which is not present in the *CellRanger* reference we used, all remaining 16 genes are detected in at least one of the samples used in the DE analysis) and chromosome Y genes (a total of 10 genes located on chromosome Y are detected in at least one sample) are differentially expressed in the expected direction (i.e. up-regulation of chromosome Y genes and down-regulation for X inactivation escape genes) for the major clusters (Figure 4d highlights these genes in the B cell cluster DE analysis). Barcode plots highlighting these genes shows clear enrichment in the major clusters, with Figure 4e showing the ranks of these sex-specific genes in the B cell cluster DE results, and Supplementary Figure S10 showing barcode plots for other clusters. Table 1 summarises the results from applying the *ROAST* gene set test (51) to the sexspecific signature across the spatial DE cluster comparisons, with statistically significant *p*-values for enrichment obtained in all cases.

**Table 1.** *ROAST p*-values from testing the enrichment of a sex-specific gene signature in the pseudo-bulk differential expression analysis between male and female spleen samples per annotated cluster or tissue region. Low count gene filtering was performed for each cluster individually, resulting in a different number of genes from the sex-specific set that could be tested for enrichment in each cluster.

| Cluster | ROAST *p*-value | Set size |
| --- | --- | --- |
| B cell | 0.000045 | 22 |
| Germinal centre | 0.00024 | 18 |
| Neutrophil | 0.00027 | 16 |
| Plasma cell | 0.00092 | 19 |
| Red pulp | 0.000015 | 22 |
| T cell | 0.00137 | 19 |

**Fig. 4.**
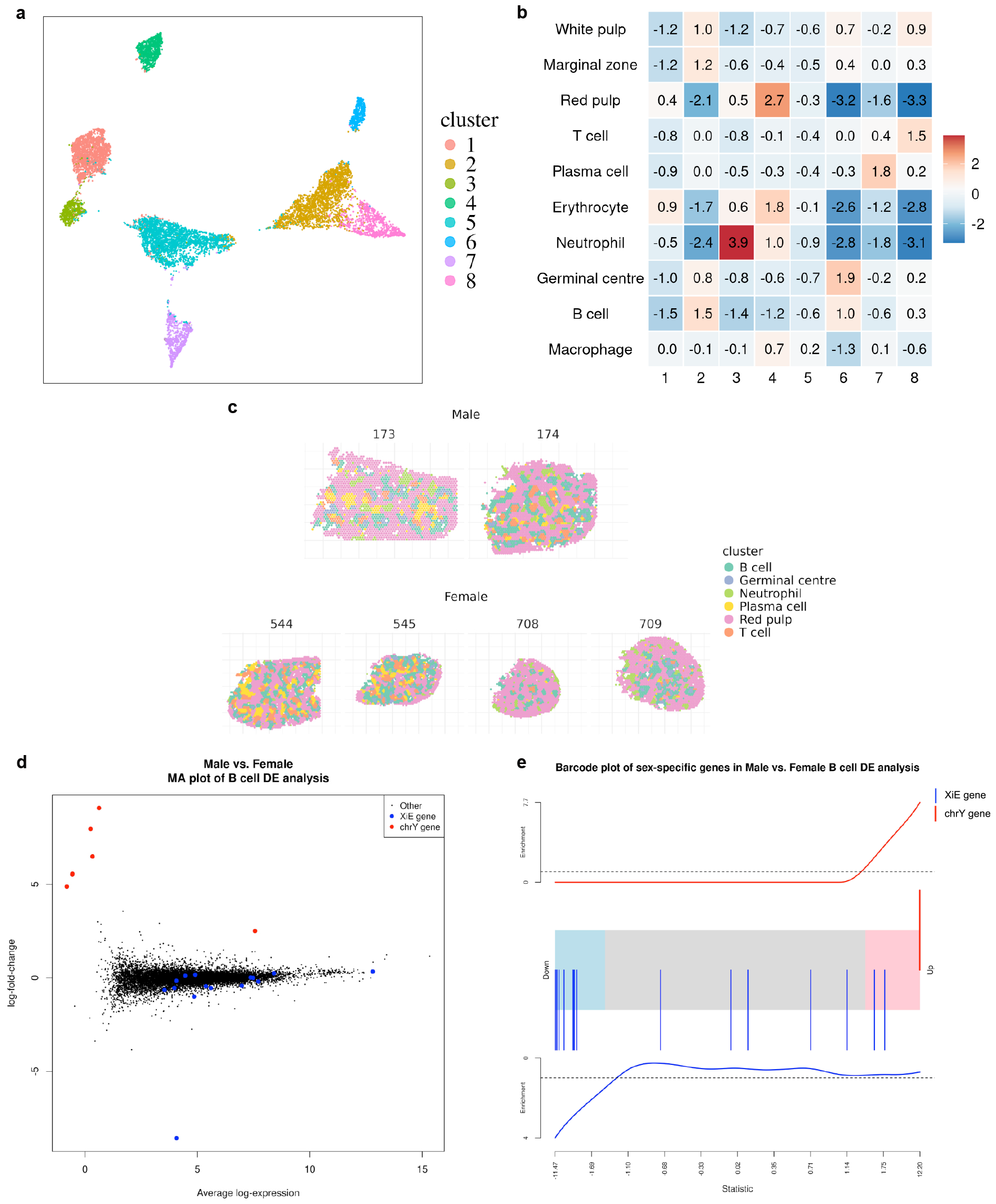
Pseudo-bulk differential expression analyses using biological sex as the ground truth. (**a**) A UMAP plot showing clusters identified by *iSC*.*MEB* across CTL (male) and WT (female) OCT samples. (**b**) Heatmap of expression scores generated using marker genes for different cell types or tissue regions expected in the spleen for each spatial cluster compared to all other clusters. (**c**) Spatial plot showing spots annotated using cluster maker gene expression from (b). (**d**) Log-fold-change vs mean expression plot of the differential expression analysis between male and female samples based on pseudo-bulk counts from cluster 2 (annotated as B cell enriched cluster). Sex-specific genes are highlighted in colour (red: chromosome Y genes, blue: genes that escape X inactivation in mouse spleen). (**e**) Barcode plot of male versus female differential expression (DE) analysis results from pseudo-bulk counts from the B cell cluster (cluster 2), with the ranks of sex-specific genes highlighted in colour (red: chromosome Y genes, blue: genes that escape X inactivation in mouse spleen).

### Comparing gene expression between replicate knockout and control spleens

We also applied pseudo-bulk DE analysis to the *T-bet* knockout (KO) and control (CTL) OCT samples and assessed the level of agreement in results with those from a previous bulk RNA-seq study, where cell sorting was used to select for germinal centre B cells from mouse spleens 15 days post-malaria infection in samples with the same genotypes (29) (see Methods). All KO and CTL spleen samples were clustered jointly and we obtained 5 distinct cell type / tissue region clusters after marker gene based annotation, including red pulp (cluster 1, 2 and 3), B cell (cluster 4), T cell (cluster 5), plasma cell (cluster 6), neutrophil (cluster 7) and germinal centre (cluster 8), which is similar to what was obtained for the previous sex-based comparison. Figure 5a shows the spatial distribution of the annotated clusters, and Supplementary Figure S11a and b shows the *iSC*.*MEB* clustering result and S11c shows the marker gene heatmap used for cluster annotation. After aggregating spot counts and filtering genes with low expression, we performed DE analysis comparing the B cell cluster from the KO samples against the CTL samples. The T-bet regulated genes from the previous study are highlighted in the spatial DE analysis results for the B cell cluster in both an MA plot (Figure 5b) and barcode plot (Figure 5c) to highlight the concordance. Enrichment of this gene set in B cell cluster was tested using *ROAST*, which gave a *p*-value of 0.005 suggesting enrichment of this signature and highlighting concordance between the Visium and bulk RNA-seq results.

**Fig. 5.**
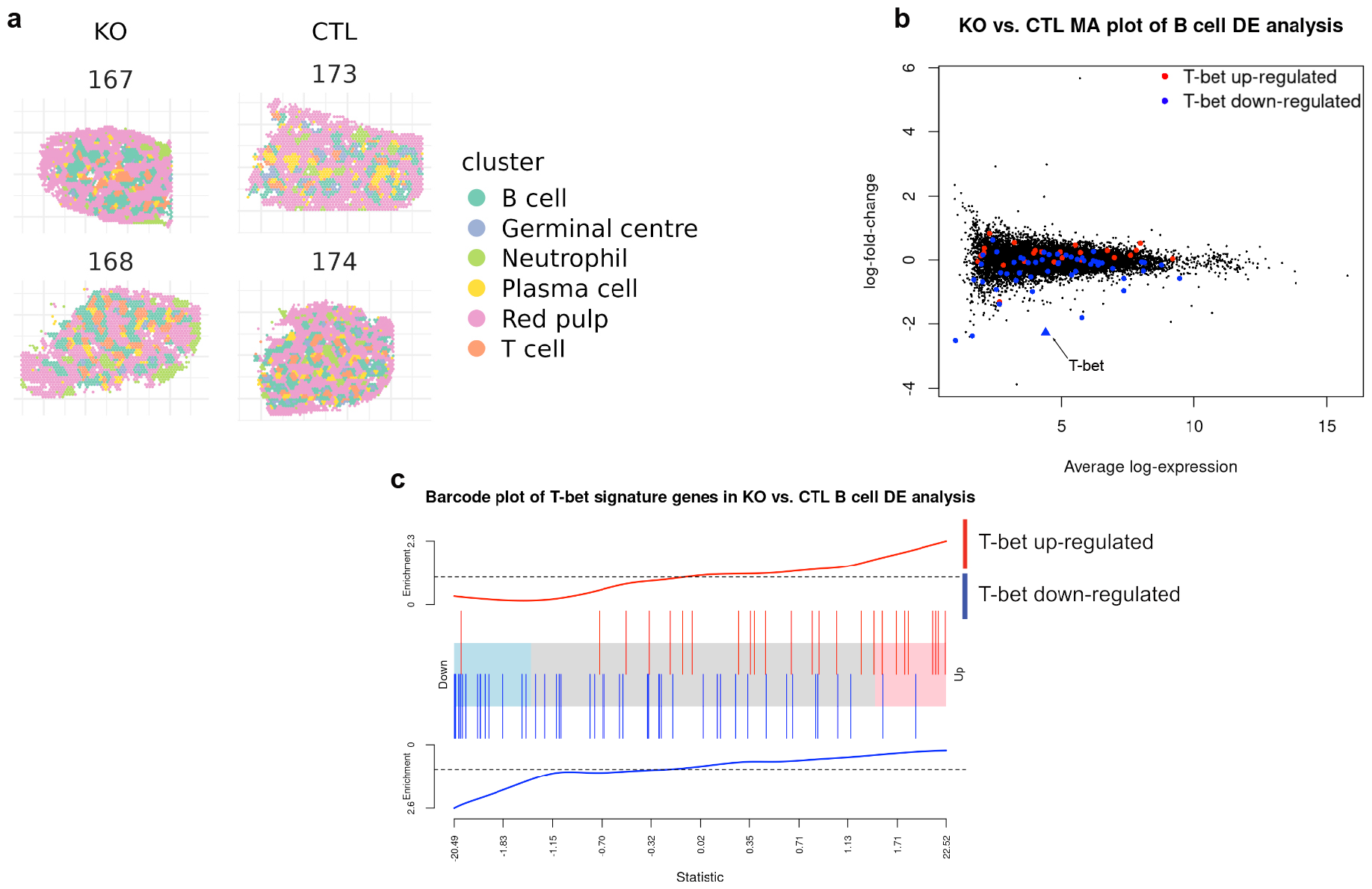
Pseudo-bulk differential expression analyses of T-bet knockouts. (**a**) Spatial plot of KO and CTL OCT samples with marker gene based annotated clusters. (**b**) Log-fold-change vs mean expression plot of the differential expression analysis between KO and CTL samples based on pseudo-bulk counts from B cell enriched cluster. A signature of differentially expressed genes in *T-bet* knockout compared to control samples from Ly *et al*. (2019) (29) are highlighted in colour (red: up-regulated genes, blue: down-regulated genes) and *T-bet* is highlighted by a blue triangle. (**c**) Barcode plot of knockout versus control differential expression analysis results from pseudo-bulk counts from the B cell cluster, with the ranks of a set of previously identified differentially expressed genes following knockout of *T-bet* highlighted in colour (red: up-regulated genes, blue: down-regulated genes).

## Discussion

We have created *SpatialBench*, a unique multi-sample spatial benchmarking dataset while profiles the mouse spleen as a reference tissue, generated using 10x Genomics’ Visium spatial technology. By harnessing Visium’s capacity to use both OCT and FFPE tissues, and integrate optimised workflows from the CytAssist platform, we systematically evaluated different sample handling protocols, data quality, and performance in various downstream analyses. We applied an effective workflow to extensively analyse the dataset, generating group-level cluster labels, identifying expected cell types, and validating existing biological knowledge of spleen dynamics during immune responses.

We observed that probe-based samples exhibited higher UMI counts, mapping confidence, and had greater proportions of retained spots after quality control than poly-A-based samples. The use of CytAssist in sample handling produced higher data quality, capturing almost all reads in spots located within tissue boundaries. This is important as only such tissue-covered reads pass the initial stage of quality control. Our findings validate 10x Genomics’ expectations for the CytAssist platform, which was designed to improve data quality through the enhanced localisation of transcripts within tissue by using optimised reagents, precise microfluidic control, and automated workflow steps. For poly-A-based samples, the observed lower mapping confidence may be attributed to the presence of unknown sequences or non-poly-adenylated transcripts in the reference, resulting in a larger proportion of unused reads. This contrasts with probe-based capture methods, where probe sets exhibit higher specificity to known sequences. In these samples, we also detected a bias showing higher UMI counts along tissue edges. A possible explanation could be sub-optimal permeabilisation conditions in our experiments, which may have impacted sensitivity and spatial resolution (52). Unlike FFPE and CytAssist workflows which use standard conditions (53, 54), OCT sample preparation involves tissue optimisation experiments during which optimal permeabilisation conditions for tissues are determined before library sequencing. Additional optimisation could be considered to mitigate this effect. It is worth noting that, however, a number of genes including some in the probe set were only detected in OCT samples, after filtering. Of these genes, there were several related to immune response and red pulp. Moreover, certain cell type marker genes were absent from the probe set. These insights may lend support to the efficacy of poly-A capture despite its poorer sensitivity.

As the field of spatial transcriptomics has evolved following single-cell transcriptomics, methods have emerged to integrate spatial information (55). Many of these were initially adapted from scRNA-seq data analyses (1) and have proven to be adaptable in procedures such as pre-processing in spatial transcriptomics workflows (56). Our own workflow incorporated scRNA-seq analysis steps following a Bioconductor workflow (36), using established packages like *scater* and *scran*. We also investigated differences between HVGs and SVGs identified with scRNA-seq and spatial analysis tools, respectively, and observed considerable overlap. This implies that the spatial distribution of cells, particularly within the major regions (red pulp and white pulp) of the spleen effectively captures biologically meaningful information in our dataset (43). From another perspective, this outcome may also suggest that HVGs suffice in downstream analyses. However, it should be noted that sensitivity is compromised in the absence of spatial information, rendering scRNA-seq tools only a short-term solution. For example, while library size is commonly considered an technical artefact and used for normalisation in single-cell analyses, variation in library sizes across tissue structures more accurately reflect biological rather than technical variation in spatial data (55). This warrants caution when adopting single-cell methods to spatial data analysis.

The efficacy of our matching scRNA-seq dataset was assessed with cell type deconvolution, showing comparable results to an established public reference (44) and outperforming it in several instances. The proportion of spots in each sample that were assigned highly confident cell type labels generated through deconvolution, served as the performance metric. Several reference-guided deconvolution methods evaluated in previous benchmarking studies (18, 21) were also explored, though we encountered various challenges in installation and computational efficiency, before advancing with *spacexr*. To streamline future analyses, we have included our scRNA-seq dataset in *SpatialBench* as a practical option, alleviating additional efforts required to source, compare, and further process different public references.

Major cell types and structures expected within a mouse spleen during malaria infection, including B cells, T cells, neutrophils, and germinal centres, were identified through a combination of various downstream processes. The spatial distributions of these components provided a detailed visualisation of the formation of germinal centres and the disrupted organisation of B and T cells, characteristic of an immune response. However, a challenge emerged concerning the labelling of red pulp, as clusters separating inner and outer regions of spots were identified. These clusters were ultimately assigned a single label based on our findings from SVG expression, deconvolution of erythrocytes, and image segmentation of red pulp and white pulp. This discrepancy could potentially be attributed to the notably lower UMI counts detected in central tissue regions compared to the edges, impacting the sensitivity of our data. The classification of cellular subtypes was less straightforward due to the limited resolution of Visium and absence of clearly defined marker gene groups beyond those for major cell types, though this could be addressed through refined iterations of deconvolution (43). Further analyses may include a more thorough examination of clusters, deconvolution, and the comparison of these results to structures identified from H&E images using object or pixel classifiers. Annotations from these images could be further used to train a custom deep learning model, which could enhance the validation of clusters and cell types.

Multi-sample spatial analysis was performed by grouping samples based on sample handling protocols, a decision that was driven by variations observed in data quality and preprocessing results. Further grouping of probe-based samples for analysis was attempted but challenges arose from persistent batch effects. *iSC*.*MEB* is one of few tools capable of processing multiple samples simultaneously (42) and allowed for the identification of concordant spatial clusters across samples within each group, enabling the generation of group-level results. Methods extended to address scenarios, where more pronounced batch effects are seen across protocols and tissue sections, would strengthen the capacity for multi-sample analysis. On a broader scale, the additional integration of high-performing spatial feature selection, clustering, and deconvolution methods would also improve workflow efficiency, offering a more streamlined approach to spatial analysis.

For differential expression analysis, a conventional workflow of clustering and aggregating spot counts to pseudo-bulk values per sample and cluster was performed. Comparisons between male and female samples recovered the expected sexspecific gene expression changes across all major cell type and region-based clusters. We also analysed DE in samples with conditional knockout of *T-bet* in mature follicular B cells, and obtained highly concordant results with a previous *T-bet* gene signature obtained from sorted germinal centre B cells from samples with the same genotypes, albeit at a slightly different time point (day 15 versus day 12). Our results highlight Visium’s potential in higher order multi-sample analyses across distinct tissue structures, while also outlining challenges in applying methods originally developed for single-cell or bulk RNA-seq experiments, especially in obtaining comparable clusters. Limited tools currently exist for multi-sample spatial analysis, and this is an area in need of further development.

We have demonstrated a comprehensive spatial preprocessing workflow and downstream analysis approach integrating SVGs and marker gene expression, cell type deconvolution, and image segmentation, both for individual samples and groups of replicate samples. Challenges in annotating some specific cell types arose potentially due to low sensitivity in several samples, and the limited resolution of Visium-generated data. However, overall our framework enabled thorough annotation of our samples, leading to detailed visualisations of the spatial context of structures within the mouse spleen. As the field of spatial transcriptomics continues to evolve, we anticipate the development of advanced methods capable of more effectively harnessing the power of spatial information in multi-sample, multi-group experiments. We envisage *SpatialBench* to be adaptable to include such methods to analyse the depth of spatial data, and to expand to more platforms in the future, such as Visium HD and others with higher spatial resolution.

## Conclusions

We present *SpatialBench*, a comprehensive Visium spatial transcriptomics dataset spanning several tissue handling protocols, that includes replicate samples and a corresponding scRNA-seq reference dataset. Our investigation into the differences between poly-A and probe-based capture library preparation protocols revealed higher quality among probe-based samples, particularly those processed with CytAssist. We also showcase the successful application of our dataset in a comprehensive analysis workflow, including steps such as pre-processing with established scRNA-seq methods and multi-sample spatial approaches to feature detection and clustering, enabling the generation of results for groups of samples. Through our analyses, we demonstrated an accurate characterisation of the cellular composition of the mouse spleen during an immune response, identifying expected cell types and structures. We anticipate that our dataset and results may serve as a practical guide to analysing data from multi-sample 10x Visium experiments.

## Methods

### Mouse spleen samples

To visualize germinal centres arising in the spleen in response to infection, a number of male *Tbx*21^*fl/fl*^*Cd*23^*Cre*^ and *Cd*23^*Cre*^ control mice as well as wild type (WT) female 8 week C57BL/6J mice were infected intravenously with 1 *×* 10^5^ *Plasmodium berghei* ANKA parasitised red blood cells (pRBCS), and then drug cured at the onset of disease symptoms using chloroquine and pyrimethamine as described previously (29). The *Tbx*21^*fl/fl*^*Cd*23^*Cre*^ conditional knockout deletes *T-bet* in mature follicular B cells.

### Sample preparation and library construction

Twelve days post-infection, mice were euthanised and spleens were fixed in 10% v/v formalin and Paraffin-Embedded (FFPE) or embedded and in Optimal Cutting Temperature compound (OCT) prior to freezing.

### Fresh frozen (OCT) Direct Placement Visium

The permeabilisation time for spleen sections was first determined using the Visium Spatial Tissue Optimization Reagents Kits User Guide. An optimal permeabilisation time of 40mins was established. 10μm sections were cut on a Cryostat and placed directly onto a Visium Spatial Gene Expression Slide. Slides were placed into slide mailers and stored at -80°C until use.

The Visium slides were processed according to the 10x Genomics Methanol Fixation, H&E Staining & Imaging for Visium Spatial Protocol followed by the fresh frozen Visium Spatial Gene Expression Reagent Kits protocol according to the manufacturer’s instructions.

### FFPE Direct Placement Visium

5μm sections were placed onto a Visium Spatial Gene Expression Slide. Slides were heated at 42°C for 3hrs on a thermocycler with a Visium PCR Adaptor then placed in a desiccator at room temperature from o/n up to 1 week. The Visium slides were processed according to the 10x Genomics Visium Spatial Gene Expression for FFPE – Deparaffinization, H&E Staining, Imaging & Decrosslinking protocol followed by the Visium Spatial Gene Expression Reagent Kits for FFPE protocol according to the manufacturer’s instructions.

### Fresh frozen (OCT) CytAssist Visium

10μm sections were cut on a Cryostat and placed directly onto a Superfrost Plus microscope slides. Slides were placed into slide mailers and stored at -80°C until use. The Visium slides were processed according to the 10x Genomics Visium CytAssist Spatial Gene Expression for Fresh Frozen – Methanol Fixation, H&E Staining, Imaging & Destaining protocol followed by the CytAssist Spatial Gene Expression Reagent Kits protocol according to the manufacturer’s instructions.

### FFPE CytAssist Visium

5μm sections were placed onto a Visium Spatial Gene Expression Slide. Slides were heated at 42°C for 3hrs on a thermocycler with a Visium PCR Adaptor then placed in a desiccator at room temperature from o/n up to 1 week. The Visium slides were processed according to the 10x Genomics Visium CytAssist Spatial Gene Expression for FFPE – Deparaffinization, H&E Staining, Imaging & Decrosslinking protocol followed by the Visium CytAssist Spatial Gene and Protein Expression Reagent Kits protocol according to the manufacturer’s instructions.

### 10x Single-cell samples

Cells were sorted on the BD Aria III (100um nozzle, 1.5ND filter) using DAPI as a live/dead cell marker. The sorted cells were centrifuged at 400xg for 5mins at 4°C and re-suspended in 25μl Cell Staining Buffer (BioLegend). 2.5μl of 1:10 TruStain FcX™ PLUS (antimouse CD16/32) (BioLegend) was added and incubated for 10mins on ice. 0.1μg mouse TotalSeq™-A HashTag and the mouse TotalSeq™-A universal cocktail v1.0 (Biolegend) diluted 1:4, were added in a total volume of 25μl, mixed and incubated on ice for 30mins. Cells were washed 3 times with Cell Staining Buffer, centrifuged at 400xg for 5mins at 4°C and resuspended in 1X PBS + 0.04% BSA. Cells were counted on the Countess III with trypan blue, pooled evenly and a total of 35,000 live cells were loaded onto a single lane of a 10x 3’ v3.1 Chip G. Gene expression libraries were produced according to the 10x Chromium Next GEM Single Cell 3’ v3.1 protocol, with HashTag and TotalSeq™-A antibody libraries produced according to the BioLegend TotalSeq™-A Antibodies and Cell Hashing protocol.

### Sequencing

All libraries were sequenced on the Illumina NextSeq2000 according to 10x guidelines.

### Pre-processing: Visium

Raw sequencing data were processed using the 10x Genomics Space Ranger 2.0.0 *mkfastq* pipeline to generate FASTQ files. Sequences were aligned to the mm10 transcriptome and gene expression counts were obtained using Space Ranger *count*. Quality metric plots were created using *ggplot2* (57) version 3.4.4 and *ggpubr* (58) version 0.6.0. Individual samples were pre-processed following a standard Bioconductor workflow for spatial transcriptomics analysis (36)

### Pre-processing: 10x Single-cell RNA-seq

Data were run through Cell Ranger 7.0.0 (59) and demultiplexing of the HTO data was performed using R/Bioconductor package *demuxmix* version 1.0.0 (60). Pre-processing was then conducted following a standard Bioconductor workflow for scRNA-seq analysis (61), using methods from *scran* (31) version 1.24.1 and *scater* (30) version 1.24.0.

### Feature selection

*nnSVG* (37) version 1.0.4 was used to first conduct feature selection. Genes were ranked by the estimated likelihood ratio value, then only those with adjusted *p*-values below 0.05 were retained. A multi-sample approach was implemented by taking the mean of gene ranks to generate lists of spatially variable genes (SVGs) for each sample type group. These were then compared with UpSet plots and venn diagrams generated using *ComplexUpset* (62, 63) version 1.3.3 and *ggvenn* (64) version 0.1.10. Highly ranked haemoglobin and immunoglobulin genes in these lists were excluded in downstream processes due to high abundance (38).

### Multi-sample cluster analysis

Pre-processed individual samples were combined into lists of *Seurat* (65) objects to create *iSC*.*MEB* objects used as inputs for multi-sample analysis with *iSC*.*MEB* (42) version 1.0. An *iSC*.*MEB* model was fitted to each object and its relevant list of SVGs, based on sample type group. Principal component analysis (PCA) was conducted on the data and the top 10 principal components (PCs) were selected for analysis. *iSC*.*MEB* then identified spatial clusters concordant across all samples in each sample type group, visualising them in a UMAP plot, and further analysed differential gene expression between clusters.

### Marker gene selection and use in cluster annotation

Marker genes for relevant cell types (Supplementary Table S3) were primarily identified from previous studies and existing literature. Specific genes for germinal centres were also chosen from a previous study (29). Zone-level marker genes for analyses comparing red pulp and white pulp were derived from individual or combined cell type marker gene lists. Some zone-specific genes such as *Hba-a1* (46) were also included here. A cluster scoring approach was developed by firstly summing the expression of groups of cell type marker genes for each spot in a cluster for each sample, then normalising by the number of spots per cluster. This calculation was repeated across all spots in all clusters and a cluster score was calculated as the log-fold-change between one cluster and all other clusters. Heatmaps of cluster scores and all spatial expression plots were created using *ggplot2* version 3.4.4.

### Cell type deconvolution

Cell type deconvolution was performed on each sample with *spacexr* (43) version 2.2.1, using either a matching scRNA-seq dataset or an external mouse spleen reference from the *Tabula Muris* compendium (44). Multi mode was used as recommended for Visium, accounting for more than two cell types per spot.

### Image segmentation

*QuPath* (66) version 0.4.3 was used for image segmentation on pathology images. A representative image was used for color deconvolution with “Estimate Stain Vectors”, which was then applied to the entire dataset. A training image with 15 patches was created by selecting 3-5 patches per image that were representative of background and white pulp and used to train a pixel classifier to predict two classes: white pulp and *Ignore. The trained classifier segmented white pulp across the dataset, and a tissue threshold generated a tissue mask. The red pulp mask was obtained by subtracting the white pulp annotation from the tissue mask. Red pulp and white pulp annotations were exported as a GeoJSON file, which was processed using R packages *sf* (67, 68) version 1.0-14 and *ggplot2* version 3.4.4.

### Pseudo-bulk differential expression analysis

Multisample clustering and marker gene based cluster annotation was performed on relevant datasets (OCT CTL and OCT WT replicate samples were analysed together in the Male versus Female comparison and OCT KO and OCT CTL replicate samples were analysed together in the KO versus CTL analysis). Next, pseudo-bulk counts were aggregated for each cluster in each sample and lowly expressed genes were filtered using *edgeR*’s (47, 48) *filterByExpr* function for each cluster individually. The *limma*-*voom* pipeline (49) was performed with *limma* (69) version 3.58.1 and *edgeR* version 4.0.16 to summarise data from replicates samples and compare different sample groups (e.g. Male versus Female or KO versus CTL). For the Male versus Female analysis, *ROAST* gene set testing (51) was applied to a sexspecific gene set in a directional way, with chromosome Y genes given a `gene.weight` of 1 and X-inactivation escape genes previously identified from mouse spleens (50) given a `gene.weight` of *−* 1. For the *T-bet* knockout versus control comparison, *ROAST* gene set testing (51) was applied to the spatial DE results using the significantly differentially expressed genes (those with an adjusted *p*-value cut-off *<* 0.05, 109 genes) from Ly *et al*. (2019) (29), where a similar knock-down and infection module was used to investigate *T-bet*’s role in germinal centres at Day 15 post malaria infection. The *t*-statistics from the previous study were used as `gene.weight`s in *ROAST* to account for both the directionality and confidence of the previous DE results.

### scRNA-seq analysis

Clustering was performed with *clusterCells* from *scran*, using *igraph*’s Louvain method (70, 71) in a *bluster* (72) shared nearest-neighbour (SNN) graph. *SingleR* (73) version 2.0.0 was used with *celldex* (73) version 1.8.0’s mouse reference datasets, *ImmGenData* and *MouseRNAseqData*, to annotate cell types. Clusters were further refined in an iterative approach of sub-clustering, re-processing and re-assigning cluster labels.

## Abbreviations

*ADT*: Antibody Derived Tags
*CA*: CytAssist
*CTL*: Control genotype
*DE*: Differential Expression
*FFPE*: Formalin-Fixed, Paraffin-Embedded
*GEM*: gel bead-in emulsion
*HTO*: HashTag oligo
*HVG*: Hyper-Variable Gene
*KO*: knockout genotype
*OCT*: Optimal Cutting Temperature compound
*PCA*: Principal Component Analysis
*RIN*: RNA Integrity Number
*scRNA-seq*: single-cell RNA-sequencing
*SVG*: Spatially Variable Gene
*WT*: Wild Type genotype.

## Declarations

### Ethics approval and consent to participate

All experiments were performed in compliance with the Walter and Eliza Hall Institute Animal Ethics Committee requirements.

## Consent for publication

Not applicable.

## Availability of data and materials

Visium and 10x scRNA-seq datasets are available from GEO under accession number GSE254652. Data analysis code is available at https://github.com/mritchielab/SpatialBench.

## Authors’ contributions

M.R.M.D and C.W. performed data analysis, generated figures and wrote the manuscript. C.W.L. and D.A.-Z. planned and supervised data collection and analysis, interpreted results and wrote the manuscript. C.J.A.A and L.L. generated data. P.F.H., C.J.S. and Y.C. contributed to study design and performed data analysis. L.J.I. provided tissue samples and contributed to study design. P.R. and R.K.H.Y. performed data analysis. K.L.R. supervised data analysis. D.S.H., R.B and M.E.R. designed the study, supervised data generation, analysis and interpretation and wrote the manuscript. All authors read and approved the final manuscript.

## Acknowledgements

We thank Catherine King, Geoffrey McDermott, Spontaneous Russell and James Fraser from 10x Genomics for feedback on our analysis and helpful insights into the technology and Kathleen Zeglinsksi for creating the *SpatialBench* project logo.

