## Supplementary Tables and Figures for "Spotlight on 10x Visium: a multi-sample protocol comparison of spatial technologies"

| Expt. | Seq. date | Sample type | KO | | CTL | | WT | | | | | | | | | Sex | Spatial protocol |
|---|---|---|---|---|---|---|---|---|---|---|---|---|---|---|---|---|---|
| 2 | 05-Dec-2022 | OCT | 167 | 168 | 173 | 174 | | | | | | | | | | M | Poly-A-based |
| 2 | 05-Dec-2022 | OCT | | | | | 544 | 545* | | | | 708* | 709 | | | F | Poly-A-based |
| 4 | 09-Mar-2023 | OCT CA | | | | | | | | | | | 709 | | 713 | F | Probe-based |
| 1 | 05-Aug-2022 | FFPE | | | | | | | 460 | 462 | 463 | | | | | F | Probe-based |
| 3 | 11-Jan-2023 | FFPE | | | | | | | | | | 708 | 709 | 710 | 713 | F | Probe-based |
| 4 | 09-Mar-2023 | FFPE CA | | | | | | | | | | | 709 | | 713 | F | Probe-based |
| | 21-Nov-22 | 10X single-cell | | | | | | | | | | 708 | 709 | | 713 | F | - |

**Table S1.** Summary of experiments, separated by OCT experiments (green) and FFPE experiments (blue). A separate single-cell experiment is included. Within the OCT/FFPE blocks, the experiments (Expt.) are ordered by date of sequencing run (Seq. date). Mouse sample IDs are grouped by genotype, where 'KO' is knockout of gene Tbet, CTL is control, and WT is wildtype. Experiment 2 was sequenced over 2 runs, on 5-Dec-22 and 3-Jan-23, to get the desired number of reads for each sample, except for samples 545 and 708 (marked by asterisks), which were only sequenced once on 5-Dec-22. The earlier date is used in the table. Experiment 4 includes 4 samples in the same run; they are presented as separate rows for OCT CA and FFPE CA.

| Expt. | Sample ID | Sample type | UMI ($n$) | UMI under tissue ($n$) | UMI removed ($n$) | UMI removed ($p$) | Median UMI count per spot under tissue ($n$) | Spots under tissue ($n$) | Spots under tissue ($p$) | Spots under tissue removed ($n$) | Spots under tissue removed ($p$) | Spots under tissue remaining ($p$) |
|---|---|---|---|---|---|---|---|---|---|---|---|---|
| 2 | 167 | OCT | 22185170 | 17150758 | 6049715 | 0.27 | 6423 | 2244 | 0.45 | 157 | 0.07 | 0.93 |
| 2 | 168 | OCT | 32584312 | 27868192 | 10659225 | 0.33 | 10084 | 2528 | 0.51 | 565 | 0.22 | 0.78 |
| 2 | 173 | OCT | 27908663 | 22230187 | 8581870 | 0.31 | 8746 | 2213 | 0.44 | 261 | 0.12 | 0.88 |
| 2 | 174 | OCT | 37333606 | 32175421 | 11840769 | 0.32 | 10491 | 2708 | 0.54 | 466 | 0.17 | 0.83 |
| 2 | 544 | OCT | 43776345 | 37876110 | 14680141 | 0.34 | 9773 | 3224 | 0.65 | 524 | 0.16 | 0.84 |
| 2 | 545 | OCT | 24807223 | 19213033 | 9826229 | 0.40 | 8672 | 2048 | 0.41 | 393 | 0.19 | 0.81 |
| 2 | 708 | OCT | 18538682 | 11853984 | 10829283 | 0.58 | 4624 | 1877 | 0.38 | 393 | 0.21 | 0.79 |
| 2 | 709 | OCT | 26473533 | 20773865 | 10862579 | 0.41 | 8062 | 2222 | 0.45 | 424 | 0.19 | 0.81 |
| 4 | 709 | OCT CA | 98876765 | 97892653 | 3269478 | 0.03 | 31796 | 2553 | 0.52 | 239 | 0.09 | 0.91 |
| 4 | 713 | OCT CA | 92074484 | 90999089 | 2156662 | 0.02 | 28311 | 2498 | 0.50 | 197 | 0.08 | 0.92 |
| 1 | 460 | FFPE | 37691995 | 24418332 | 13366678 | 0.35 | 34715 | 592 | 0.12 | 9 | 0.02 | 0.98 |
| 1 | 462 | FFPE | 38542861 | 27562657 | 11117985 | 0.29 | 36142 | 637 | 0.13 | 10 | 0.02 | 0.98 |
| 1 | 463 | FFPE | 37489533 | 27084993 | 10730631 | 0.29 | 29313 | 833 | 0.17 | 70 | 0.08 | 0.92 |
| 3 | 708 | FFPE | 63569576 | 46950218 | 17248292 | 0.27 | 18958 | 2034 | 0.41 | 267 | 0.13 | 0.87 |
| 3 | 709 | FFPE | 66361852 | 55123330 | 12079037 | 0.18 | 23426 | 1910 | 0.38 | 87 | 0.05 | 0.95 |
| 3 | 710 | FFPE | 61183497 | 50875755 | 10762044 | 0.18 | 21558 | 1922 | 0.39 | 56 | 0.03 | 0.97 |
| 3 | 713 | FFPE | 61381430 | 48288502 | 13344269 | 0.22 | 22979 | 1640 | 0.33 | 34 | 0.02 | 0.98 |
| 4 | 709 | FFPE CA | 71097828 | 69786672 | 2089205 | 0.03 | 27624 | 2002 | 0.40 | 68 | 0.03 | 0.97 |
| 4 | 713 | FFPE CA | 25474425 | 24585379 | 941141 | 0.04 | 11488 | 1711 | 0.34 | 15 | 0.01 | 0.99 |

**Table S2.** Summary of UMI counts and spots, separated by OCT experiments (green) and FFPE experiments (blue). Numbers ($n$) and proportions ($p$) of UMI counts per spot, valid UMI counts under tissue, and spots under tissue, before and after quality control, are reported. Experiment (Expt.) 4 includes 4 samples in the same run; they are presented as separate rows for OCT CA and FFPE CA.

| Cell type/splenic region | Selected marker genes |
|---|---|
| B cell | *Cd19, Cd22, Ighd, Cd5* |
| T cell | *Trac, Cd3d, Cd4, Cd3e, Cd8a* |
| Macrophage | *Cd274, Marco, Csf1r, Adgre1, Cd209b, Cd206, Cd80, Mac1, Cd68* |
| Neutrophil | *S100a9, S100a8, Ngp* |
| Erythrocyte | *Car2, Car1, Klf1* |
| Plasma cell | *Cd38, Cd138, Xbp1, Irf4, Prdm1, Cd27, Cd319, Mum1* |
| Germinal centre | *Cxcr4, Cd83, Bcl6, Rgs13, Aicda* |
| Marginal zone | *Marco, Lyz2, Ighd, Igfbp7, Igfbp3, Ly6d* |
| Red pulp | *Ifitm3, C1qc, Hmox1, Hba-a1, Klf1* |
| White pulp | *Ighd, Cd19, Trac, Trbc2* |

**Table S3.** A list of marker genes for cell types or different tissue regions expected in the mouse spleen selected from previous studies and existing literature. These marker genes were used for cluster scoring of spatial clusters obtained from *iSC.MEB* to inform their annotation.

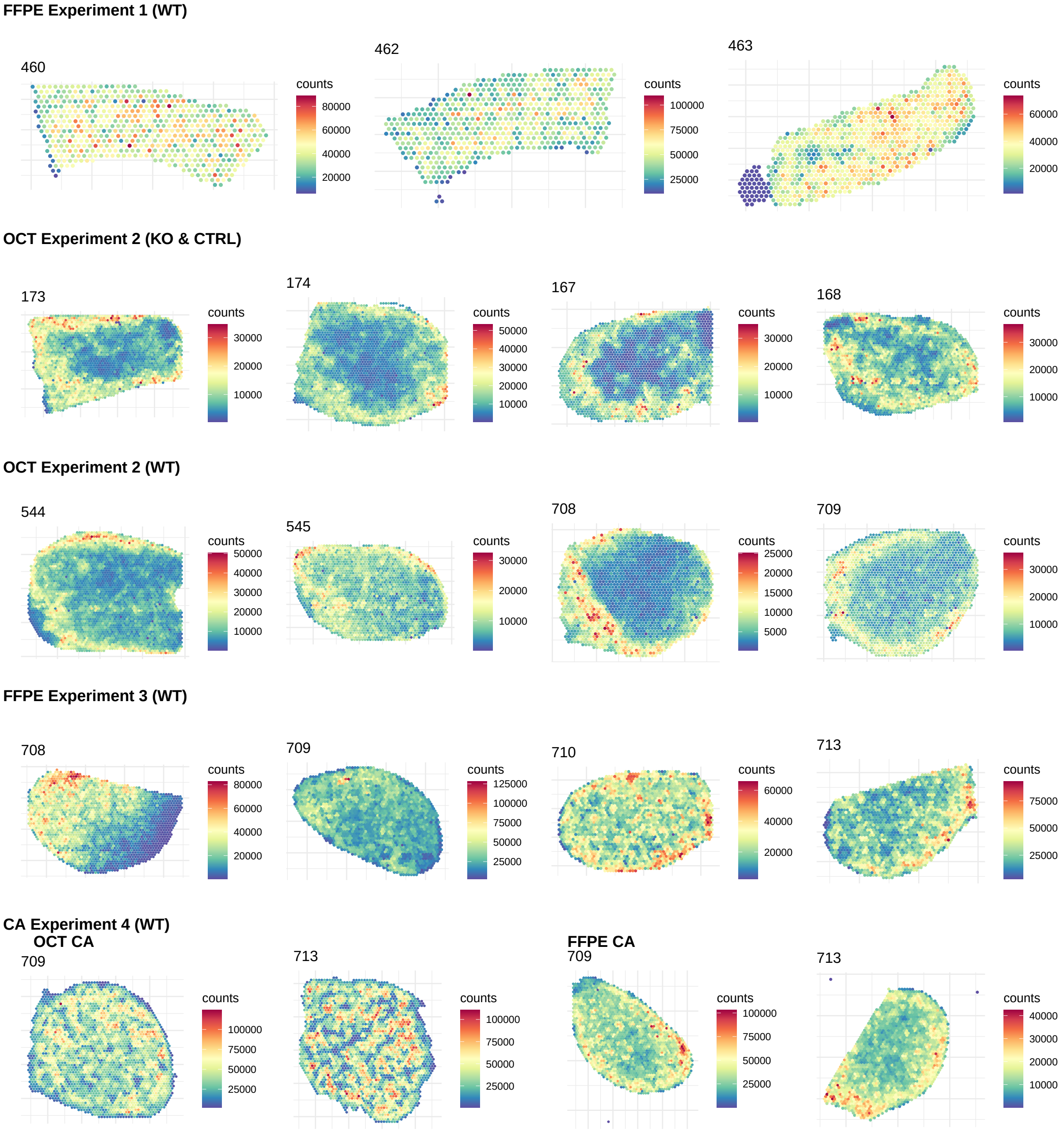

**Figure S1.** Plots showing the spatial distribution of UMI counts for all samples in the study, grouped by sample type.

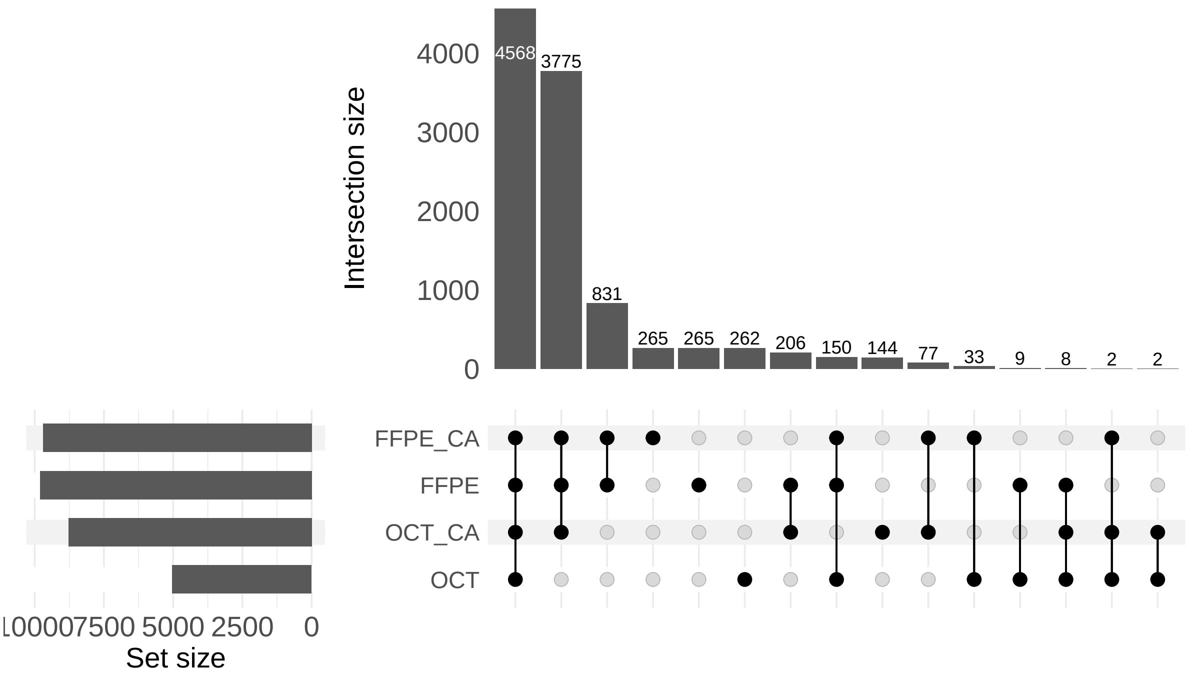

**Figure S2.** Upset plot showing the overlap of detected genes in all WT samples, categorised by sample type. Detected genes are defined as genes with a count of 3 or more in at least 1% of spots.

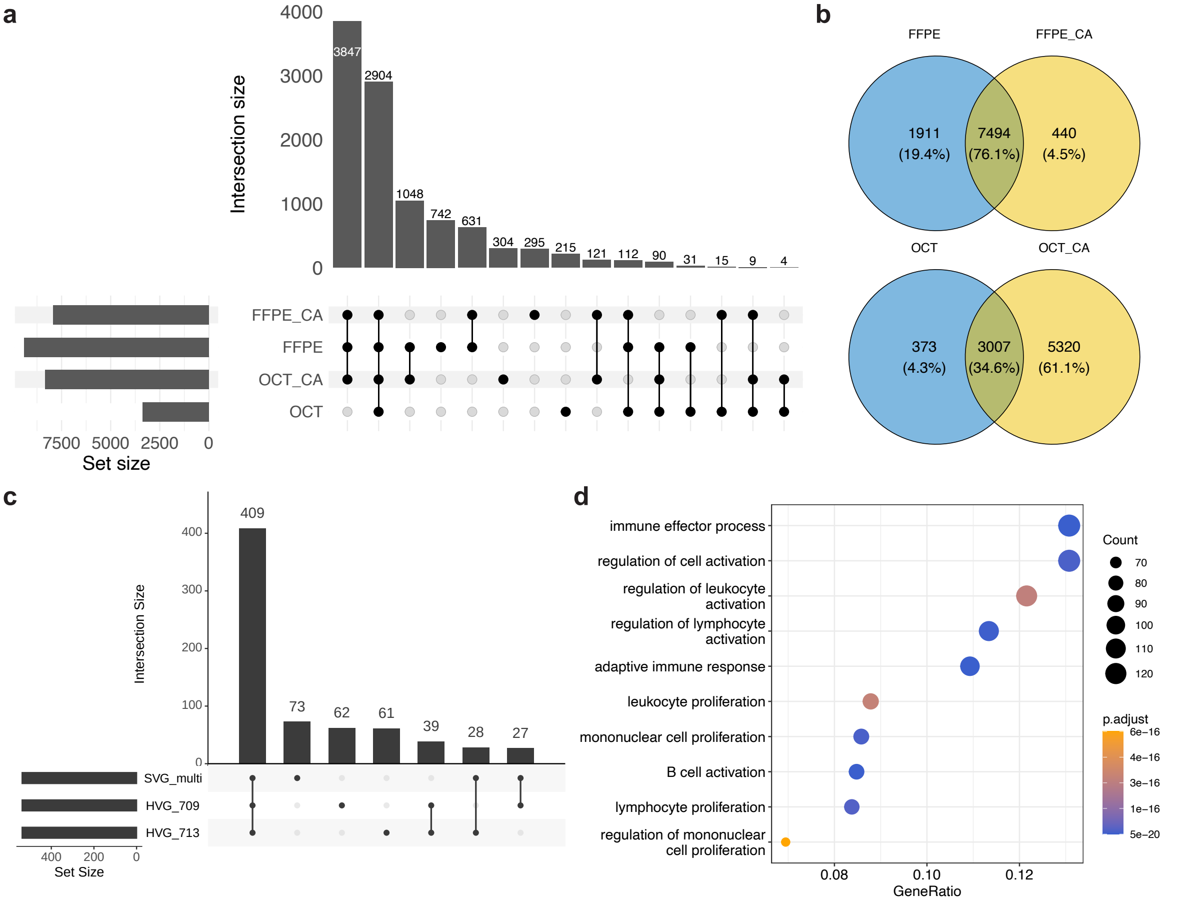

**Figure S3.** Analysis of SVGs across all sample types and HVGs in FFPE CA samples. (**a**) Upset plot of the overlap of all SVGs in each sample type for WT samples only. (**b**) Venn diagrams showing unique and overlapping SVGs between FFPE samples and between OCT samples, with and without CytAssist, among all SVGs. (**c**) Upset plot showing how variable genes intersect in FFPE CA samples, at the point of highest overlap between all gene lists (537 genes). (**d**) Heatmap showing the most highly enriched Gene Ontology (GO) terms of the top 1,000 SVGs in FFPE CA samples. 'GeneRatio' denotes the ratio of top SVGs to the total number of SVGs for a particular GO term. An adjusted $p$-value cutoff of 0.05 was used.

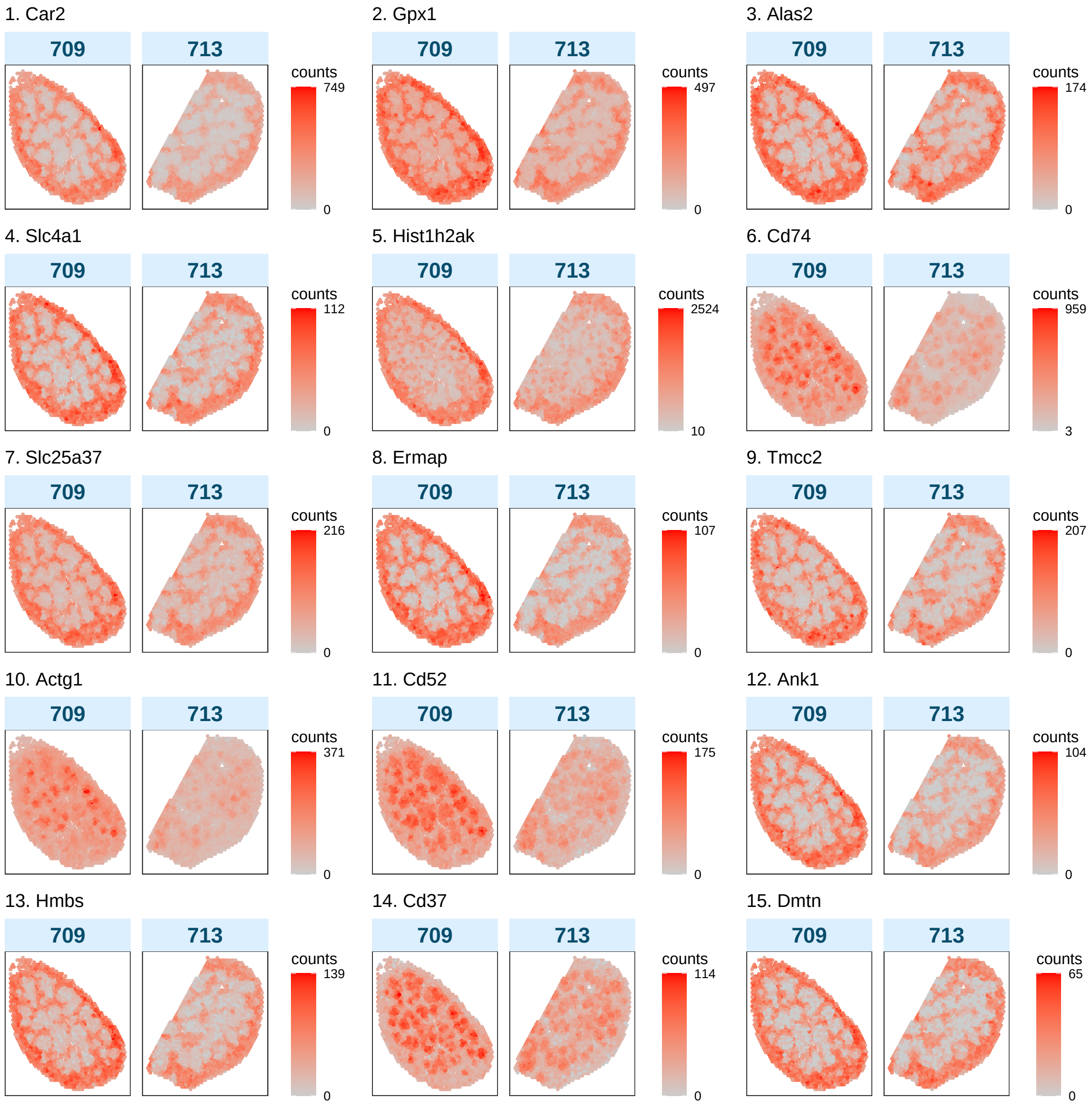

**Figure S4.** Plots displaying the spatial expression of the top 15 SVGs in FFPE CA samples, identified by *nnSVG*. Heatmaps display UMI counts per spot.

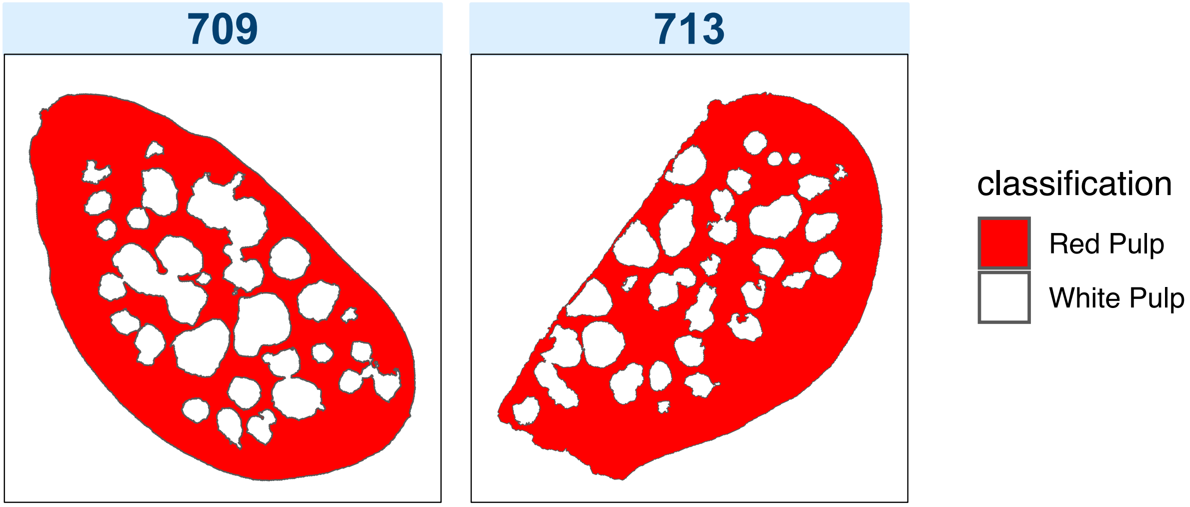

**Figure S5.** Plots showing predicted red pulp and white pulp regions in FFPE CA samples. Annotations were generated by a classifier trained with a series of pathology images imported into *QuPath* version 0.4.3.

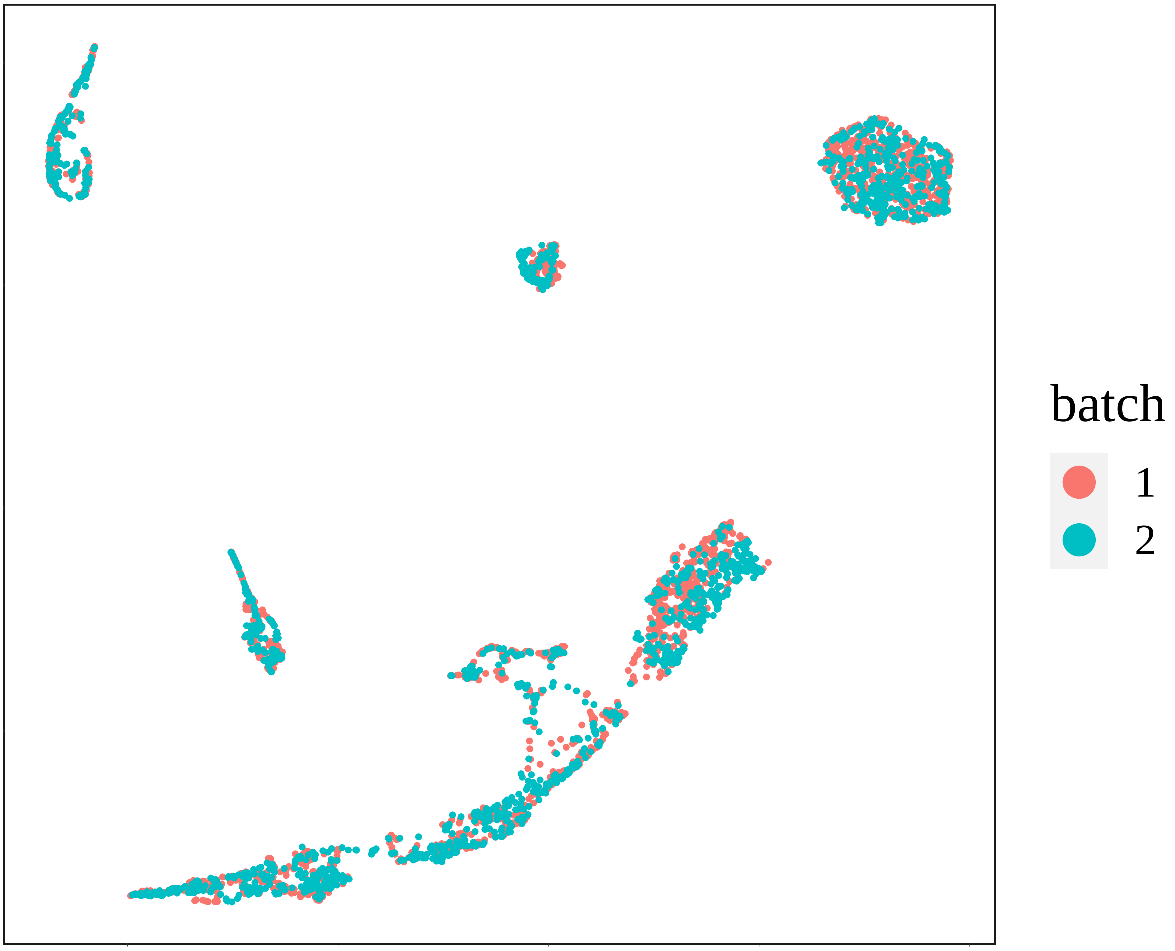

**Figure S6.** A UMAP plot generated with the *iSC.MEB* package showing the spatial distribution of FFPE CA sample batches, with 'batch' referring to the samples where 1 is sample 709 and 2 is sample 713.

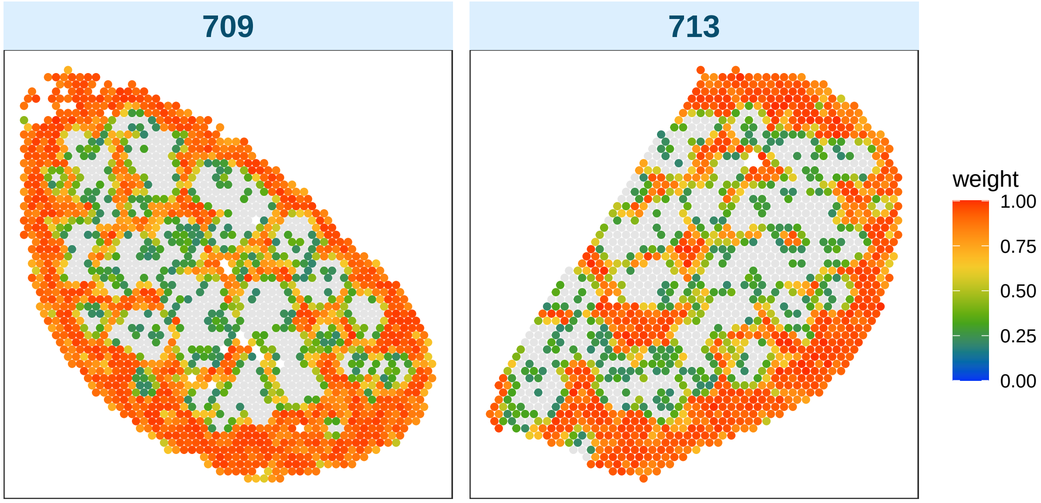

**Figure S7.** Plots showing the spatial distribution of normalised confident weights, predicting erythrocyte proportions, in each spot. Weights were generated by *spacexr* version 2.2.1 during cell type deconvolution of FFPE CA samples.

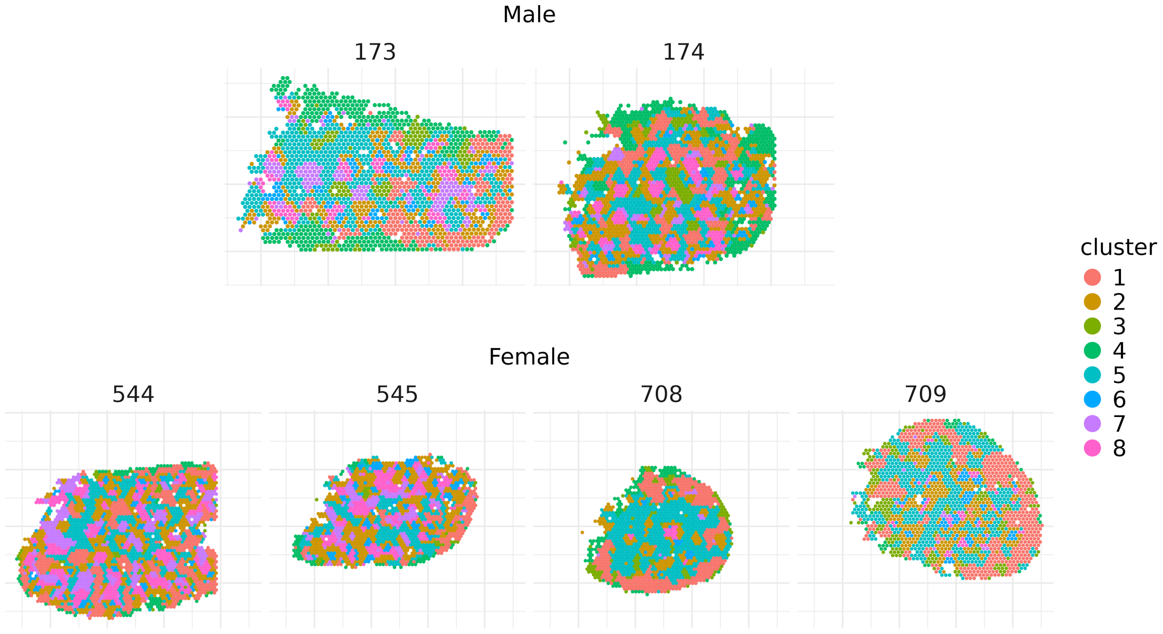

**Figure S8.** Spatial plot of *iSC.MEB* clustering result for each WT and CTL OCT spleen sample. Male samples are shown in the top row, and female samples in the bottom row.

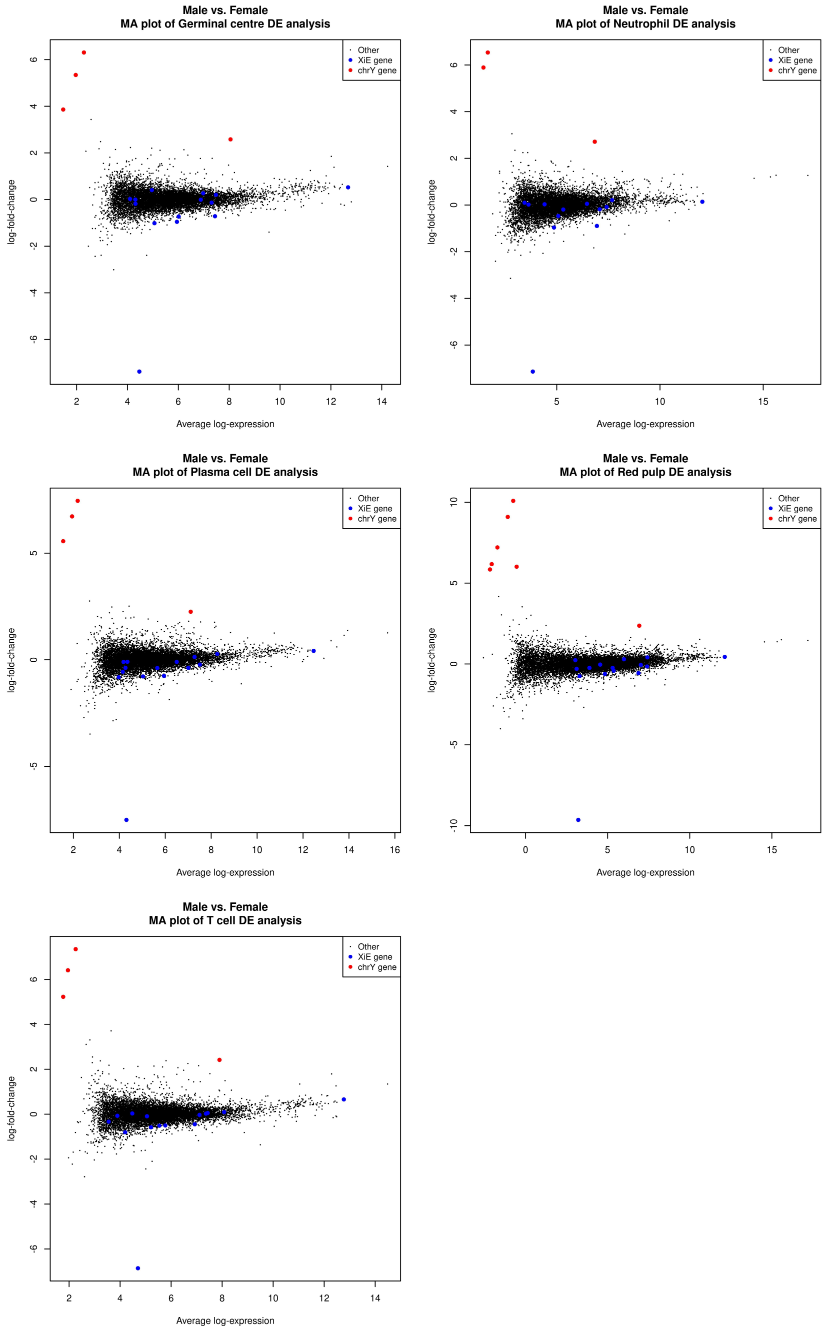

**Figure S9.** Log-fold-change vs mean expression (MA) plots of the differential expression analysis between male and female samples based on pseudo-bulk counts for different clusters. Sex-specific genes are highlighted in colour (blue: genes that escape X inactivation in mouse spleen, red: chromosome Y genes).

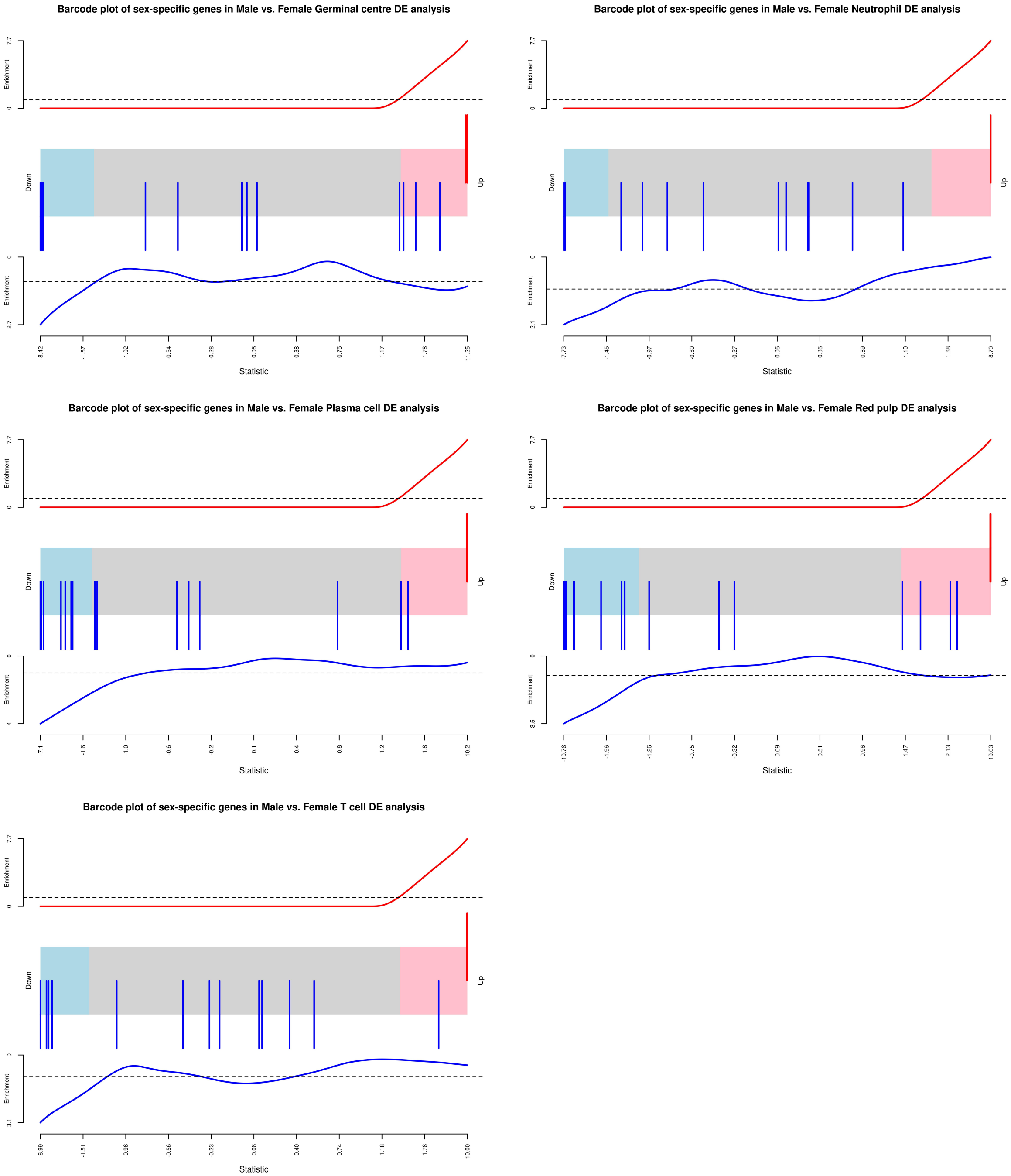

**Figure S10.** Barcode plot of male vs female differential expression analysis results from pseudo-bulk counts for each cluster, with the ranks of sex-specific signature genes highlighted in colour (blue: genes that escape X inactivation in mouse spleen, red: chromosome Y genes). *ROAST* enrichment $p$-values for this signature for each cluster are available in Table 1.

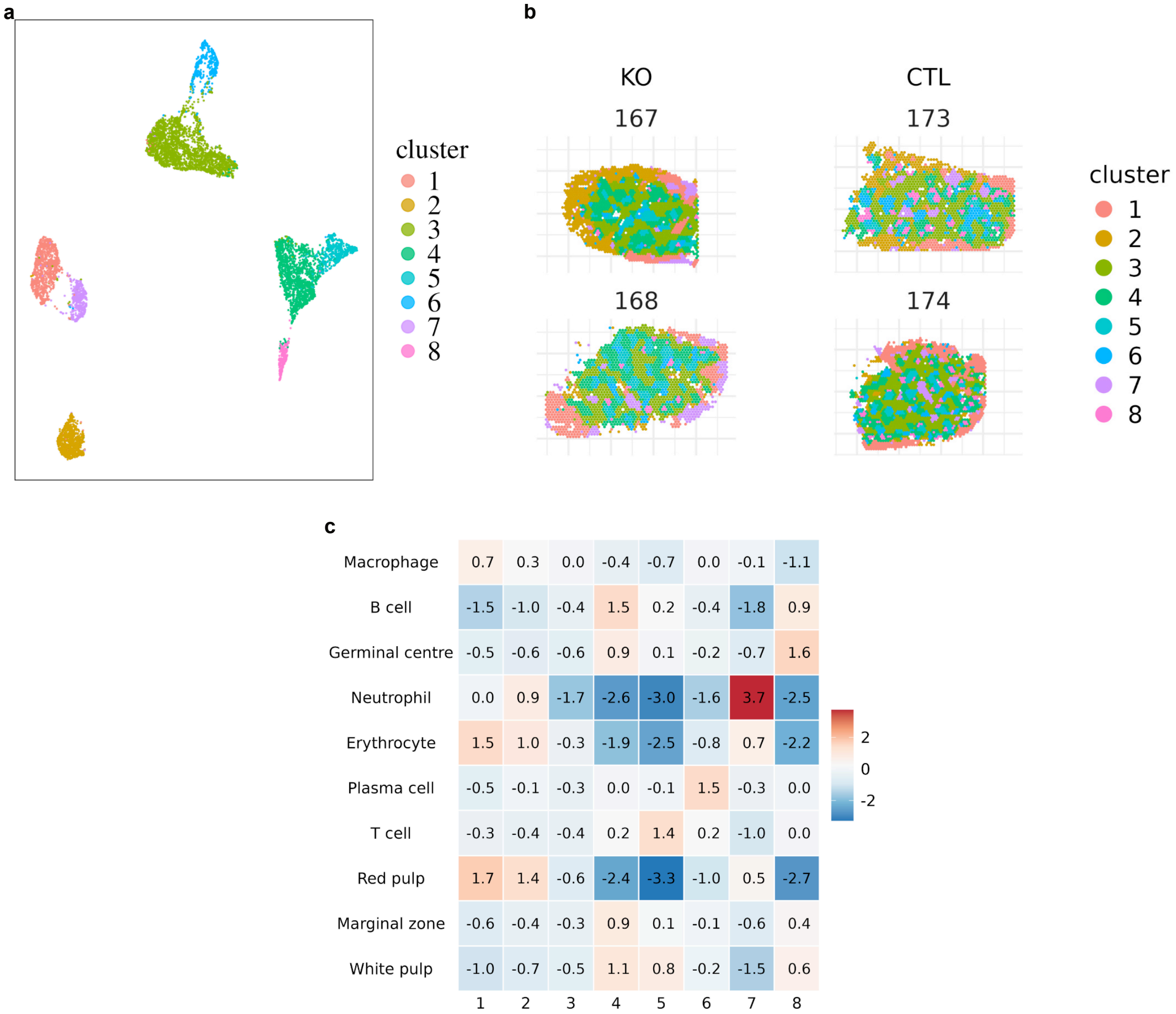

**Figure S11.** (**a**) UMAP of *iSC.MEB* clustering result for the KO and CTL OCT samples. (**b**) Spatial plot of *iSC.MEB* clustering result for KO and CTL samples, with KO samples displayed on the left and CTL samples on the right. (**c**) Heatmap of expression scores generated using marker genes for different cell types or tissue regions expected in the spleen for each spatial cluster compared to all other clusters.
